# Phosphaphenalene Gold(I) Complexes as Broad-Spectrum Antivirals Against Emerging Flaviruses

**DOI:** 10.64898/2026.02.10.705012

**Authors:** Pablo Fuentes-Soriano, Blanca Palmero-Casanova, Laura Albentosa-González, Antonio Mas, Carlos Romero-Nieto, Rosario Sabariegos

## Abstract

Flaviviruses are emerging and re-emerging pathogens of major global health concern that can replicate in hosts of different phyla, including humans. Among them, Zika virus (ZIKV) is associated with severe neurological outcomes, including neonatal microcephaly, congenital malformations, and Guillain-Barré syndrome in adults. Usutu virus (USUV), another mosquito-borne flavivirus, primarily infects birds and has been linked to neurological symptoms in humans. Bagaza virus (BAGV) significantly affects the wildlife of some bird species and carries a noteworthy risk of zoonotic transmission. Despite their clinical significance, approved antiviral options for flaviviral infections (including ZIKV, USUV, and BAGV) are not available, and current management focuses primarily on providing symptomatic relief and supportive care. To address this therapeutic gap, we evaluated the antiviral activity of five phosphaphenalene-based gold (I) complexes. These phosphaphenalene-derived compounds exhibit straightforward and versatile chemistry, allowing access to derivatives with improved stability, bioactivity, selectivity, and cytotoxicity, thereby enhancing their biological activities. Two compounds showed potent inhibition of ZIKV, USUV, and BAGV replication at low micromolar concentrations, with ZIKV displaying greater sensitivity. ZIKV titers were reduced by up to three orders of magnitude in a dose-dependent fashion. Mechanistic studies revealed that both compounds inhibited thioredoxin reductase and disrupted autophagy. This study represents the first demonstration of the potential of phosphaphenalene-based gold (I) complexes as promising candidates against flavivirus infections, offering a promising foundation for the development of urgently needed antiviral therapies with potential impact on global health.

## INTRODUCTION

Vector-borne viral infections represent an increasing global health threat. Among these, orthoflaviviruses have gained particular attention because of their rapid geographic spread and the absence of effective antiviral treatments(1). *Orthoflavivirus* genus comprises positive-sense ssRNA viruses, including major pathogens such as dengue virus (DENV), Zika virus (ZIKV), West Nile virus (WNV), Usutu virus (USUV), and Bagaza virus (BAGV)(1-5). Environmental and socioeconomic changes have expanded mosquito vector habitats, driving the global emergence and re-emergence of flavivirus outbreaks, with over 400 million people infected with flaviviruses annually(6, 7). Current challenges include understanding flavivirus biology and developing effective vaccines and antivirals. This study addresses both aspects by focusing on ZIKV, USUV, and BAGV biology and antiviral development.

ZIKV and USUV pose a growing public health threat(1, 2). The first human case of ZIKV was described in 1964(8), and since then, its spread has raised global concern because of its neurological complications and congenital effects(7, 9-13). USUV was first isolated in Africa in 1959 but was not introduced in Europe until 1996(14). It has been associated with neurological complications and encephalitis in birds and humans(14-16). BAGV, first isolated in the Central African Republic in 1966, circulates between mosquitoes and birds(17). Human infection was suggested by neutralizing antibodies detected during a 1966 encephalitis outbreak in India(18) and codon usage adaptation(17), suggesting possible cross-species transmission.

Despite their severe impact on public health, no specific antiviral treatment is currently available(19, 20). In this context, gold(I) complexes have attracted significant attention as potential antimicrobial agents because of their unique chemical properties and ability to interact with various host cell proteins essential for microbial replication(21, 22). Thus, gold(I) complexes, typically comprising a gold atom coordinated with phosphines or thiolates, can disrupt critical cellular processes and have demonstrated antiviral activity against a wide range of viruses, including retroviruses (HIV), coronaviruses (SARS-CoV-2), and arboviruses such as Chikungunya and ZIKV(23-31). This broad spectrum of antiviral activity marks gold(I) complexes as promising candidates for antiviral drug development.

Several design strategies have been employed to improve the stability, bioactivity, and selectivity of gold(I) complexes. While traditional approaches have focused on trisubstituted phosphine ligands, heterocyclic phosphorus ligands, such as phosphaphenalenes, offer a fundamentally different and highly promising alternative(32-34). Unlike classical phosphines, phosphaphenalenes possess a fused six-membered phosphorus heterocycle, which imparts unique electronic and structural properties(34). Despite the infancy of their chemistry, these ligands have been demonstrated to effectively coordinate with gold(I) and, owing to their extended π-conjugation and rigid backbone, give rise to distinct coordination geometries and stronger metal–ligand interactions. Additionally, phosphaphenalenes possess versatile post-functionalization reactions, enabling precise fine-tuning of solubility, lipophilicity, and reactivity, which are crucial for improving bioavailability and reducing cytotoxicity(34). This chemical versatility renders phosphaphenalenes a powerful platform for the design and development of next-generation gold-based therapeutics.

Herein, we report a new family of phosphaphenalene-gold(I) complexes with antiviral activity against flavivirus and provide evidences consistent with the inhibition of thioredoxin reductase (TrxR) activity and autophagy perturbation. Thus, beyond providing new molecular scaffolds for therapeutic exploration, this study offers key insights into the design principles of gold(I)-based molecules as highly efficient antiviral agents.

## MATERIALS AND METHODS

### Compound synthesis

Reactions were carried out in dry glassware under an inert atmosphere of purified nitrogen using Schlenk techniques. All solvents (THF, Toluene, DCM) were used directly from the solvent purification system MB SPS-800. Lithium diisopropylamine, P,P-diphenylchlorophosphine, trichlorosilane, chloro(dimethylsulfide)gold(I), and ethyl xanthogenate were purchased from commercial suppliers.

### NMR and mass spectrometry

^1^H, ^13^C, and ^31^P NMR spectra, as well as COSY spectra, were recorded on a 400 MHz Varian Inova NMR spectrometer. Chemical shifts were expressed as parts per million (ppm, δ) and referenced to external 85% H_3_PO_4_ (^31^P) or solvent signals (^1^H/^13^C): CDCl_3_ (7.27/77.16 ppm) and CD_2_Cl_2_ (5.33/53.80 ppm), as internal standards. Signal descriptions included s = singlet, d = doublet, t = triplet, q = quartet, m = multiplet, and br = broad. All coupling constants are absolute values, and J values are expressed in Hertz (Hz).

HR-ESI spectra were measured using the NUCLEUS analytical service (University of Salamanca, Spain). GC-MS was performed using a 7250 GC/Q-TOF (Agilent Technologies).

### Virus strains, cell lines and cell viability

ZIKV strain PRV-ABC59 (ATCC VR-1843) and USUV strain 939/01 (Austria, 2001) were propagated as previously described(35). BAGV was isolated from an infected animal(36). Vero cells were maintained in DMEM (Sigma-Aldrich) supplemented with 5% FBS (Sigma-Aldrich), penicillin–streptomycin (Sigma-Aldrich), and 1 mM HEPES (Gibco, The Netherlands) at 37°C with 5% CO₂.

Cell viability (CC₅₀) was determined by seeding 1×10⁴ Vero cells per well in 96-well plates and treating them with increasing drug concentrations for 48 h, as described previously(35).

### Infection and virus titration

Infection experiments were performed as previously described(35). Vero cells (1×10⁵) were seeded in 24-well plates and infected 24 h later with ZIKV, USUV, or BAGV at the indicated m.o.i. Cells were treated with the indicated drug concentrations and incubated for 48 h. Supernatants were collected for viral titration. Mock-infected cells were used as controls.

Viral titers were determined by TCID₅₀ assay as previously described(35). Briefly, 1×10⁴ Vero cells were seeded in 96-well plates and infected with 10-fold serial dilutions of supernatants. Cytopathic effects were evaluated 4–5 days post-infection.

### Inhibitory Concentration (IC_50_) assays

Vero cells (1x10^5^) were infected with ZIKV (m.o.i.=0.5), USUV (m.o.i.=0.5) or BAGV (m.o.i.=1). Cells were treated at concentrations below CC₅₀ and incubated for 48 h at 37°C in 5% CO₂. Supernatants were collected, and viral titers determined as described above. The selectivity index (SI) was calculated as CC₅₀/IC₅₀.

### RNA extraction and RT-PCR analysis

ZIKV RNA was extracted from 100 µL of virus–drug supernatant or from infected cells (24-well plates) 48 h post-infection using the GeneJET RNA Purification Kit (Thermo Fisher Scientific). Semi-quantitative RT-PCR was performed using MMLV reverse transcriptase and DreamTaq polymerase (Thermo Fisher Scientific) with specific primers (F:5’-cagatggaccttgcaaggttcc-3’, R:5’-cttgaaacagaggagatcccgcagataccatc-3’) yielding ∼700 bp amplicons. Products were resolved on 1% agarose gels and quantified (Quantity One, Bio-Rad).

For RT-qPCR, ZIKV and USUV RNA were extracted as described above. Viral genomes were quantified using NZYSupreme One-step RT-qPCR Probe Master Mix on a CFX Duet Real-Time PCR System (Bio-Rad), as previously described(37, 38).

### Western blot

Vero cells (5×10⁵) were seeded into 24-well plates, treated for 48 h, washed with PBS, lysed, and total protein was quantified as previously described(35). Equal amounts of protein (50 µg) were resolved by SDS-PAGE (12% for LC3-I/II; 10% for SQSTM1/p62) and transferred for immunoblotting, as previously described(35, 37). LC3-I and LC3-II were detected using α-LC3 (Rabbit, PA1-16931, Invitrogen), SQSTM1/p62 with α-p62 (Rabbit, PA5-20839, Invitrogen), and GAPDH with α-GAPDH (Mouse, GT239 HRP-conjugated, GeneTex). HRP-conjugated goat anti-rabbit IgG (31460, Thermo Fisher Scientific) was used as secondary antibody.

### Thioredoxin reductase (TrxR) activity

Thioredoxin reductase (TrxR) activity was measured using a Thioredoxin Reductase Colorimetric Assay Kit (Cayman, Michigan, #1007892) following the manufacturer’s instructions. TrxR activity in treated cells was expressed as percentage relative to untreated controls, normalized to 100%.

### Statistical analysis

Data are expressed as mean ± SD and analyzed using GraphPad Prism v8. Comparisons were performed using Student’s *t*-test or ANOVA, followed by Tukey’s post hoc test. Statistical significance is indicated as \**p* < 0.05, \*\**p* < 0.01, \*\*\**p* < 0.001.

## RESULTS

### Viral titers and viral replication inhibition

To screen the bioactivity of the phosphaphenalene gold(I) complexes, we first used compounds **1** – **4**(33) (Figure 1). All these gold(I) complexes contain a phosphaphenalene core with a fused thiophene and differ in the nature of the anionic moiety attached to the gold atom. This allowed us to systematically investigate the effects of these moieties on the antiviral activity of the complexes. Compound **1** (**C1**) contains a chloride atom, compound **2** (**C2**) a xanthate fragment, compound **3** (**C3**) a thiocyanate group, and compound **4** (**C4**) a thiosugar derivative (Figure 1A). These moieties have been found to lead to distinct bioactivities in previous studies(33). Data on the structures of these compounds are provided in the Supplementary Information. To determine the concentration range of each compound for subsequent infection and treatment experiments, we first determined the CC_50_ value for each compound, as described in the Materials and Methods section. The values obtained ranged from 1.1 µM for **C4** to 6.7 µM for **C2** (Table 1).

**FIGURE 1.**
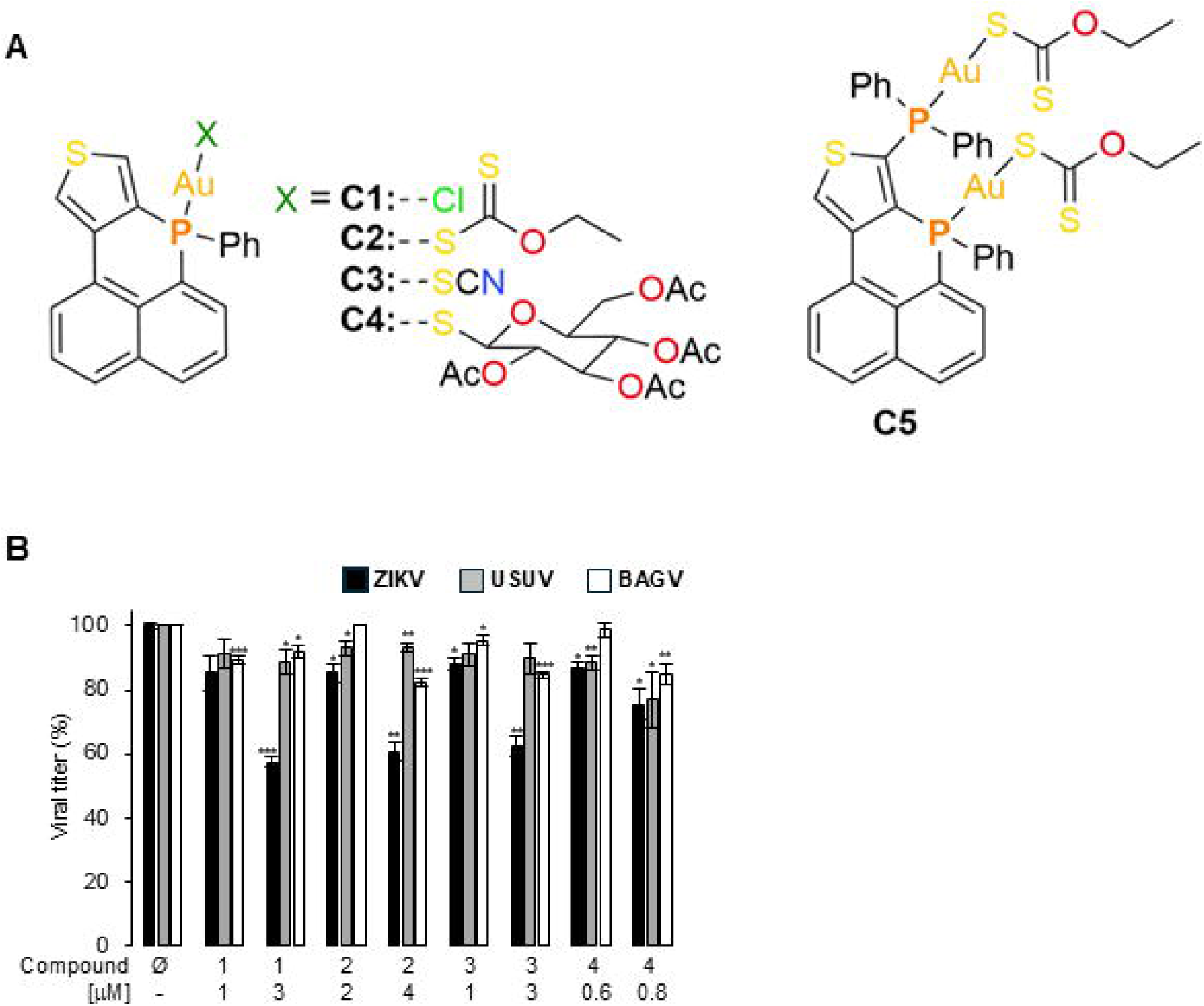
Structure and effect of phosphaphenalene derivatives C1 to C5 on ZIKV, USUV, and BAGV replication. **A.** Core molecule and the different ligands (X) of phosphaphenalene compounds **1** - **5**. **B.** Effect of compounds **1** - **4** on replication of ZIKV, USUV and BAGV. Vero cells were infected at a m.o.i. of 0.5 and treated with or without compounds **1** to **4** for 48 h at the indicated concentrations. Virus titers were obtained from untreated control (C) and treated cells. Data shown are the average ± SD of at least two independent biological experiments performed in triplicate and normalized against control cells. Student’s t-test was used for statistical analysis, where significance of differences is represented as follows: ns, no statistical difference; * p<0,05; ** p<0.01; *** p<0.001.

**Table 1.**
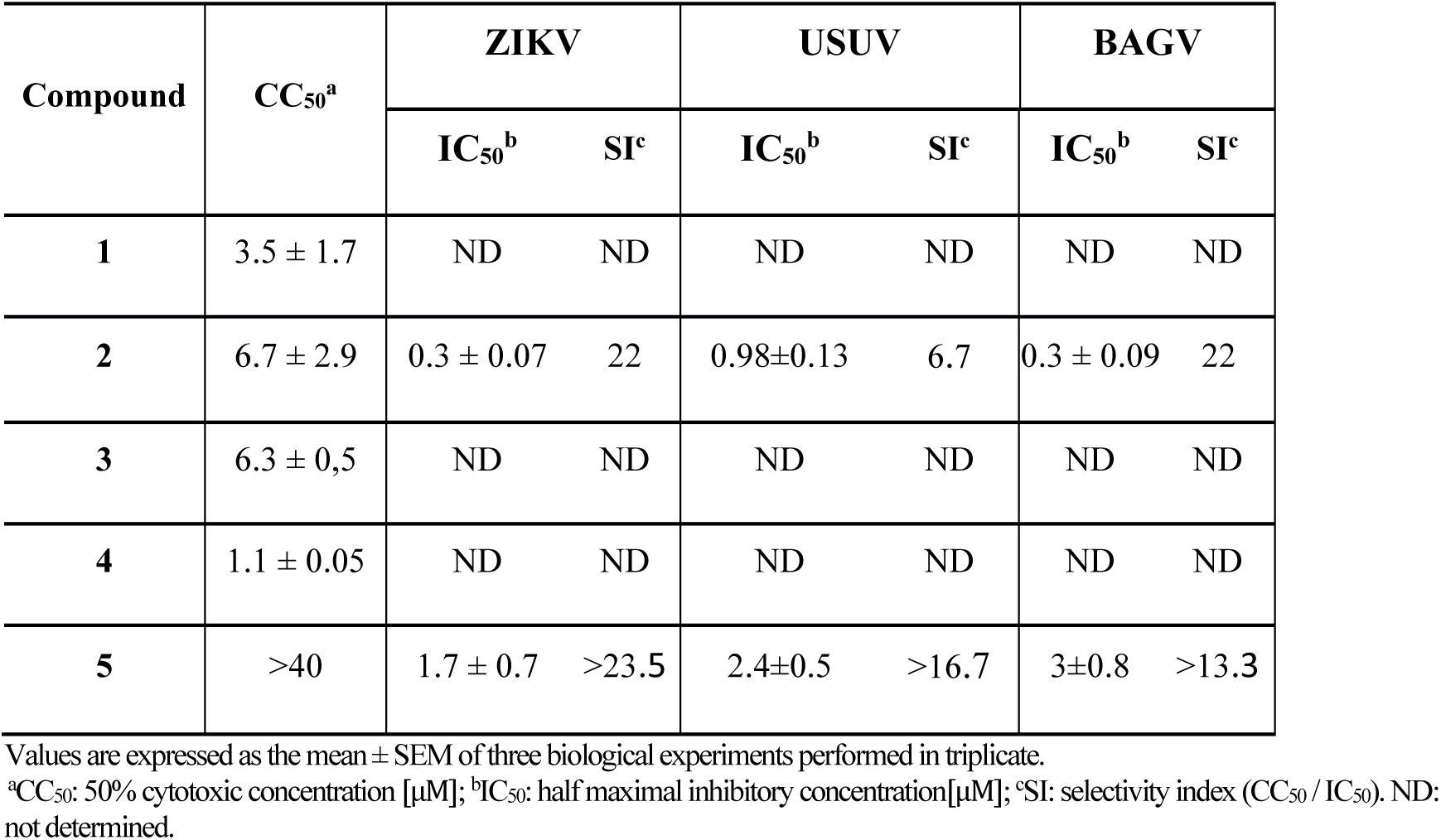
CC50, IC50 and SI of compounds 1 to 5 at 48 h.

To evaluate whether **C1**–**C4** inhibited ZIKV, USUV, and BAGV replication in cell culture, virus-infected Vero cells were treated with two different drug concentrations (Figure 1B). The concentrations chosen were lower than the CC_50_ values of each compound and ensured cell viability above 85% (Table 1). All compounds exhibited similar antiviral activities at the concentrations used. Therefore, we decided to continue with **C2** as the starting point for chemical modifications because it exhibited the lowest cytotoxicity.

Next, we quantified effectiveness of **C2** against ZIKV, USUV, and BAGV by measuring viral titers across increasing drug concentrations. As shown in Figure 2A, the ZIKV titer was reduced by three logs (from 6.1 to 3.1) at the highest **C2** concentration. USUV titers decreased by 1.3 logs (Figure 2B), whereas BAGV titers decreased by 1.8 logs (Figure 2C). These results clearly demonstrate that **C2** exhibits antiviral activity against these viruses, with a stronger effect on ZIKV. These data were used to calculate the corresponding IC_50_ values, which were all on a low-micromolar scale: 0.3±0.07 (ZIKV), 0.98±0.13 (USUV), and 0.3±0.09 (BAGV) (Table 1). Using these values, we calculated the selective index (SI), which was 22 for ZIKV and BAGV and 6.7 for USUV (Table 1).

**FIGURE 2.**
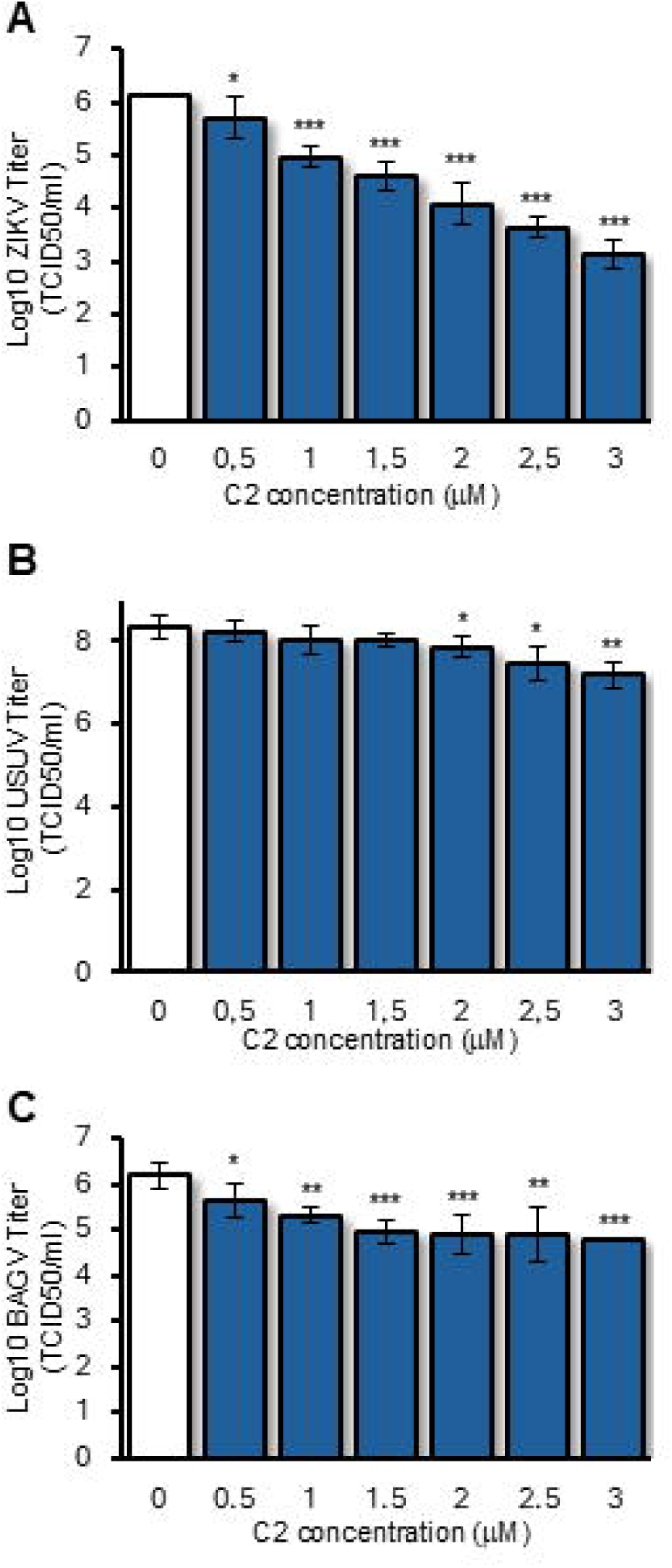
Effect of compound 2 on the replication of ZIKV, USUV, and BAGV. Vero cells were infected with ZIKV (**A**), USUV (**B**), and BAGV (**C**) at an m.o.i. of 0.5, and treated with **C2** at the indicated concentrations for 48 h. Virus titers were obtained from untreated (0) and treated cells. Data are presented as the mean ± SD of at least two independent biological experiments performed in triplicate. Statistical analysis was performed as described in Figure 1.

### Synthesis of a new phosphaphenalene derivative and biological properties

**C2**, which possesses an anionic xanthate group, exhibited the lowest toxicity (CC_50_ = 6.7 µM, Table 1). Therefore, phosphaphenalene-gold(I) derivatives with xanthate moieties appear to be the best candidates for further investigation. Based on these data, we synthesized compound **5** (**C5**) with two active gold-xanthate moieties to maintain or enhance the antiviral activity of **C2** and further improve its cellular toxicity (Figure 1A). The synthesis is shown in Figure 3. We first tested the possibility of selectively functionalizing one of the alpha positions of the thiophene ring of phosphaphenalene oxide **6**. The treatment of compound **6** with lithium diisopropylamide (LDA) at low temperature, followed by a reaction with deuterated water as a reference reaction, confirmed the possibility of quantitatively installing electrophiles at position 1 of thiophene-based phosphaphenalenes. Notably, no side reactions or decomposition were observed, emphasizing the synthetic versatility of phosphaphenalenes. Thus, compound **7** (Figure 3), with two pentavalent phosphorus centers, was obtained by treating **6** with LDA, followed by a reaction with Ph_2_PCl as the electrophile and final oxidation with H_2_O_2_. To obtain complex **C9** with two gold atoms (Figure 3), the phosphorus atoms of **C7** were reduced with HSiCl_3_ and reacted with Me_2_SAuCl. The anion interchange of both chloride atoms by the two xanthates yielded novel **C5** (see Supplementary Information for details).

**FIGURE 3.**
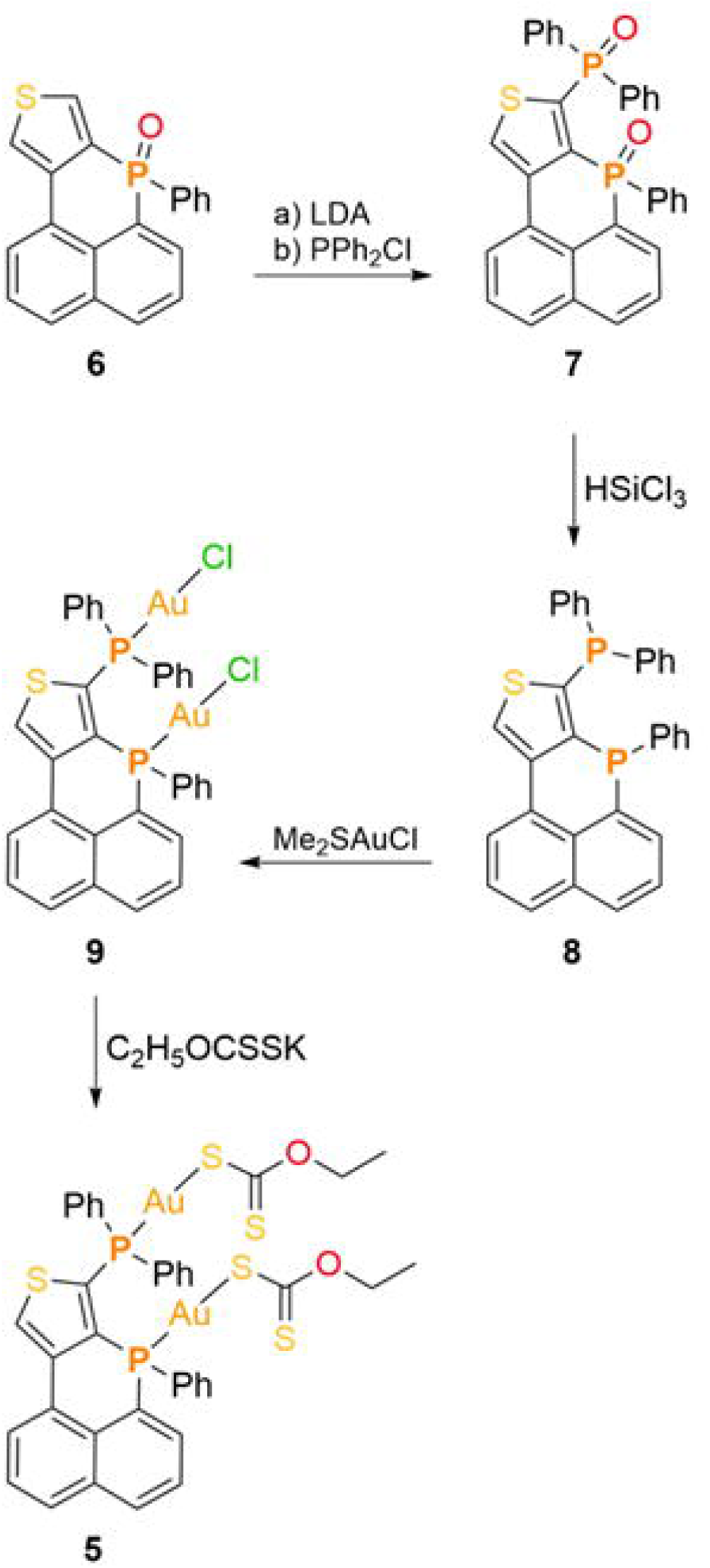
Synthetic pathway for the preparation of phosphaphenalene derivatives. The synthesis of compounds **7**, **8**, **9**, and **5,** with compound **6** as the starting material, is shown. The treatments for each chemical reaction are indicated. The chemical reactions are described in detail in the text.

### Antiviral activity of compound 5

**C5** cytotoxicity was assessed by treating cells with increasing concentrations and measuring viability, yielding a CC₅₀ above 200 µM (Table 1). Although we did not observe a decrease in cell viability, this compound tended to aggregate at concentrations above 40 µM, resulting in homogeneous monolayers on the cells. Therefore, **C5** treatment was performed at concentrations below 40 µM.

ZIKV titer was reduced by 1.7 logs (from 5.9 to 4.2) at the highest concentration of **C5** (Figure 4A). The reduction in USUV viral titer under these conditions was 1.4 logs (from 8.2 to 6,8; Figure 4B), and the reduction in BAGV vital titer was 1.4 logs (from 6.1 to 4.7; Figure 4C). Using the data from the experiments shown in Figure 4, we calculated the IC_50_ value of **C5** for each virus. The values obtained were 1.7±0.7 µM, 2.4±0.5 µM, and 3±0.8 µM for ZIKV, USUV, and BAGV, respectively (Table 1). The IC_50_ and CC_50_ values (Table 1) were used to calculate the SI. The CC_50_ value for **C5** could not be calculated, and we decided to use 40 µM to calculate the SI for this compound. The SI values were greater than 23, 16.9, and 13 for ZIKV, USUV, and BAGV, respectively. These results indicate that **C5** inhibits the replication of these flaviviruses (Figure 4), with an almost complete absence of cytotoxicity (Table 1).

**FIGURE 4.**
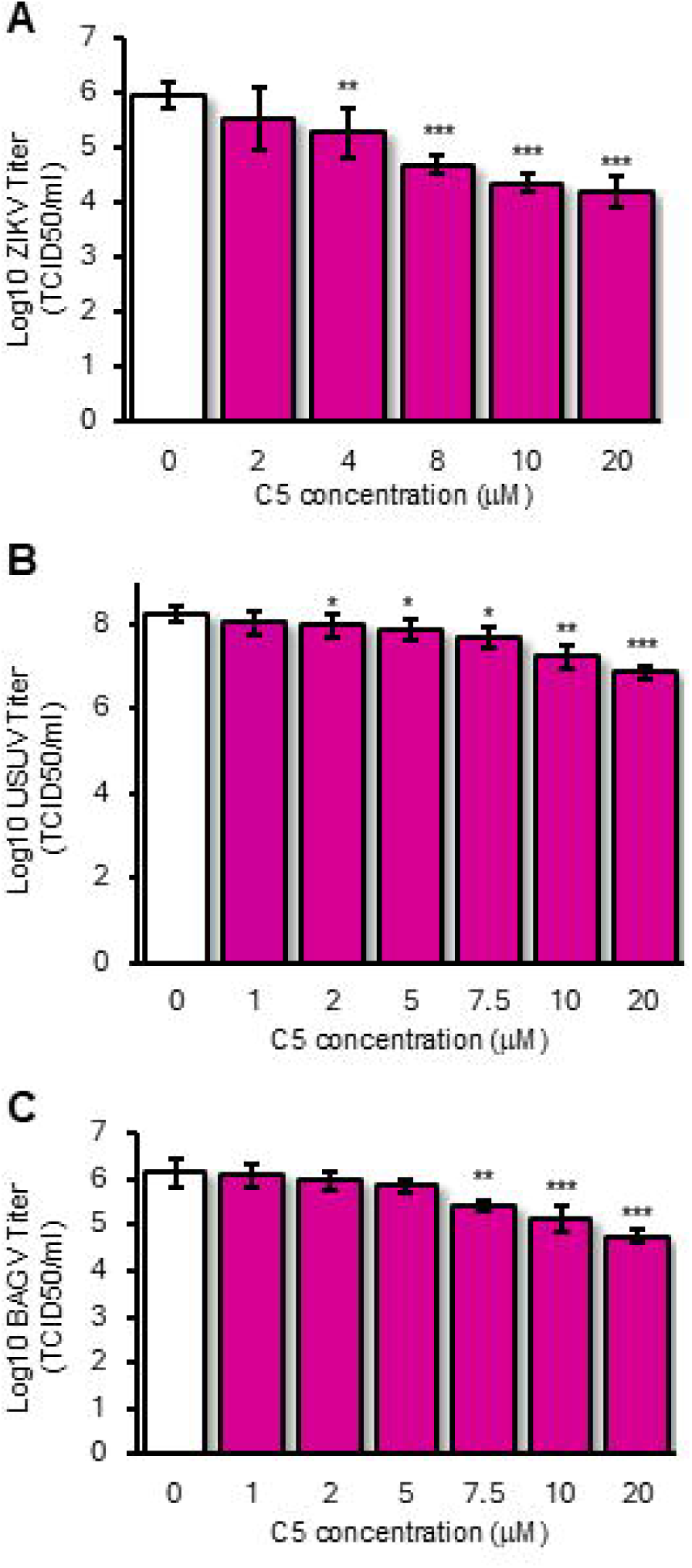
Effect of compound 5 on the replication of ZIKV, USUV, and BAGV. Vero cells were infected with ZIKV (**A**), USUV (**B**), and BAGV (**C**) at a m.o.i. of 0.5 and treated with **C5** at the indicated concentrations for 48 h. Viral titers were obtained from untreated (0) and treated cells. Data shown are the average ± SD of at least two independent biological experiments carried out in triplicate each one. We used the Student’s t-test for statistical analysis. Statistically significant differences are represented as follows: ns (no statistical significance); * p<0.05; ** p<0.01; ***p<0.001.

### Genome Quantification

To confirm the observed decrease in ZIKV titer, RT-PCR amplification assays were performed to compare semi-quantitatively the effect of treatment with these compounds on the viral genome copy number. Treatment with both **C2** and **C5** decreased the ZIKV genome copy number, as deduced from amplicon intensity (Figure 5A and 5B), consistent with viral titer reductions (Figures 2 and 4) and comparable to ribavirin (400 µM) treatment.

**FIGURE 5.**
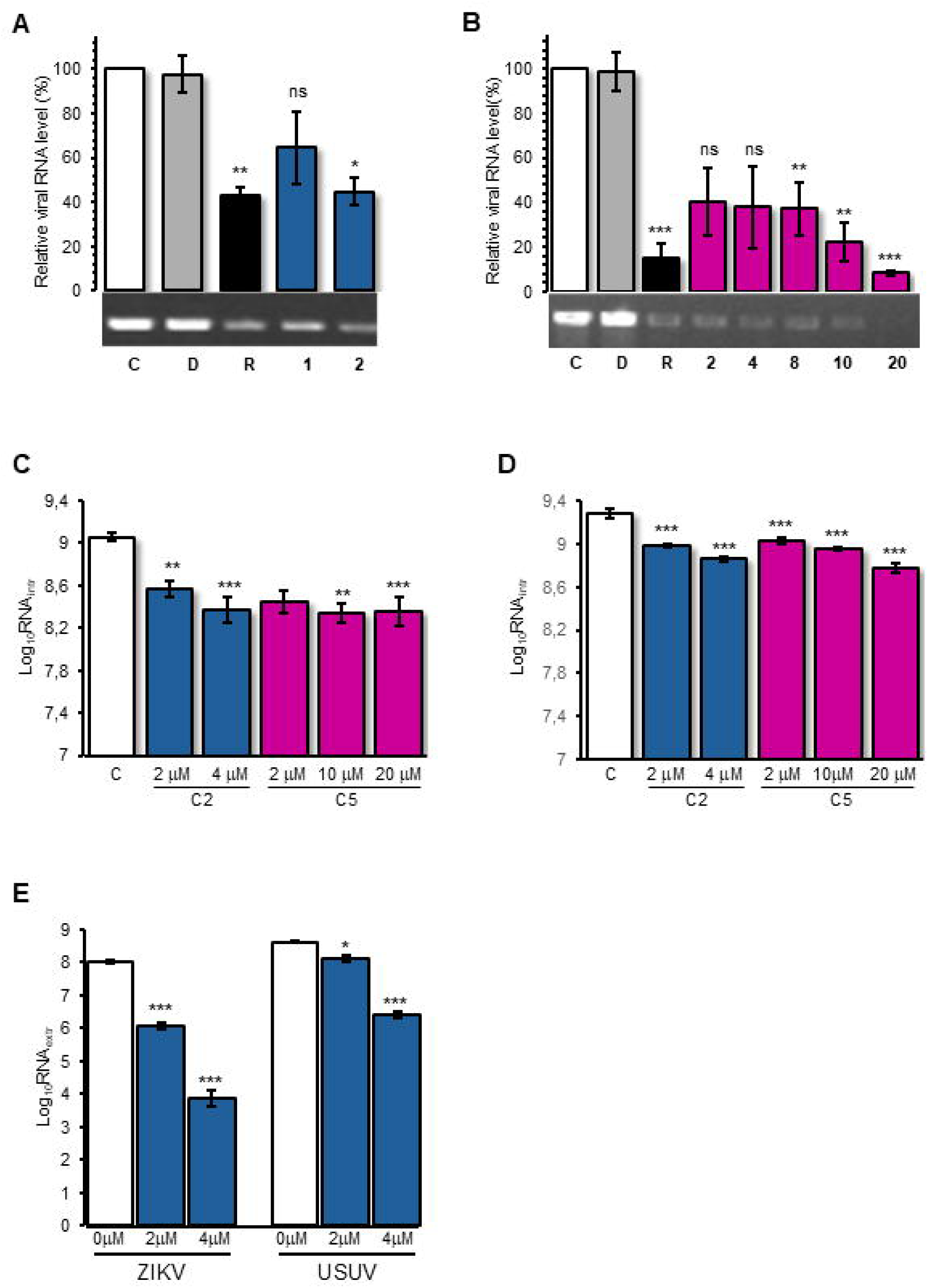
Viral genome quantification. **A** and **B**. RT-PCR assays were performed by amplifying a region of the ZIKV genome using total RNA extracted from the supernatants of cultures with ZIKV-infected cells treated with different concentrations (µM) of compounds **2** (**A**) and **5** (**B**). Untreated ZIKV-infected cells ("C") and ZIKV-infected cells treated with DMSO ("D") were used as negative controls. ZIKV-infected cells treated with ribavirin (R, 400 μM) were used as positive controls. RT-PCR products were resolved in 1% agarose gel and quantified. Quantification is presented as the mean ± SEM of three independent experiments. Student’s t-test was used for statistical analysis. **C** and **D**. Intracellular quantification of viral genomic RNA. Total RNA was extracted from cells infected with ZIKV (**C**) or USUV (**D**) and treated as indicated in the figure. **E.** Extracellular quantification of viral genomic RNA. Total RNA was extracted from cells infected with ZIKV or USUV and treated with different concentrations (µM) of **C2**. Viral genomic RNA was quantified by RT-qPCR as described in the materials and methods section. The mean ± SEM of the copy number (Log_10_) obtained from at least three independent experiments is shown. ANOVA with an ad-hoc Tukey test was used for statistical analysis. Statistically significant differences are represented as follows: ns, no statistical significance; * p<0,05; ** p<0,01., *** p<0,001.

We further quantified viral genome copies inside the cells and in the supernatants after ZIKV and USUV infection and treatment. We performed intracellular RNA quantification for **C2** and **C5** treatments using qRT-PCR, as described in the Materials and Methods section. The results (Figure 5C and 5D) showed a decrease in the viral intracellular genome RNA copy number, although to a lesser degree than the viral titer results (Figures 2 and 4). To assess the release of newly formed viral particles into the extracellular environment, we quantified extracellular viral RNA in the presence of **C2** (Figure 5E). The results showed a marked decrease in the release of newly formed viral particles. This reduction was comparable to that observed in viral titration assays (Figures 2 and 4).

### Thioredoxin reductase activity and autophagy

Gold(I) complexes exhibit a strong affinity for thiol and selenol groups, allowing them to selectively interact with proteins such as TrxR. Gold (I)-derived compounds, such as auranofin, are important for human health and have been associated with TrxR inhibition and increased autophagy(39-41). Additionally, flaviviruses have been reported to exploit autophagy pathways to enhance their replication(42). Given this background, we wondered whether the mechanistic activity of compounds **C2** and **C5** might be related to TrxR inhibition and/or alteration of the autophagic state of the cells. Thus, TrxR enzymatic activity was measured in cellular extracts treated with **C2** and **C5** and compared with that in untreated cells. The presence of both compounds led to a statistically significant reduction in TrxR activity, which was more pronounced for **C2** than for **C5** (Figure 6A), consistent with the data on inhibition of viral replication (Figures 2 and 4).

**FIGURE 6.**
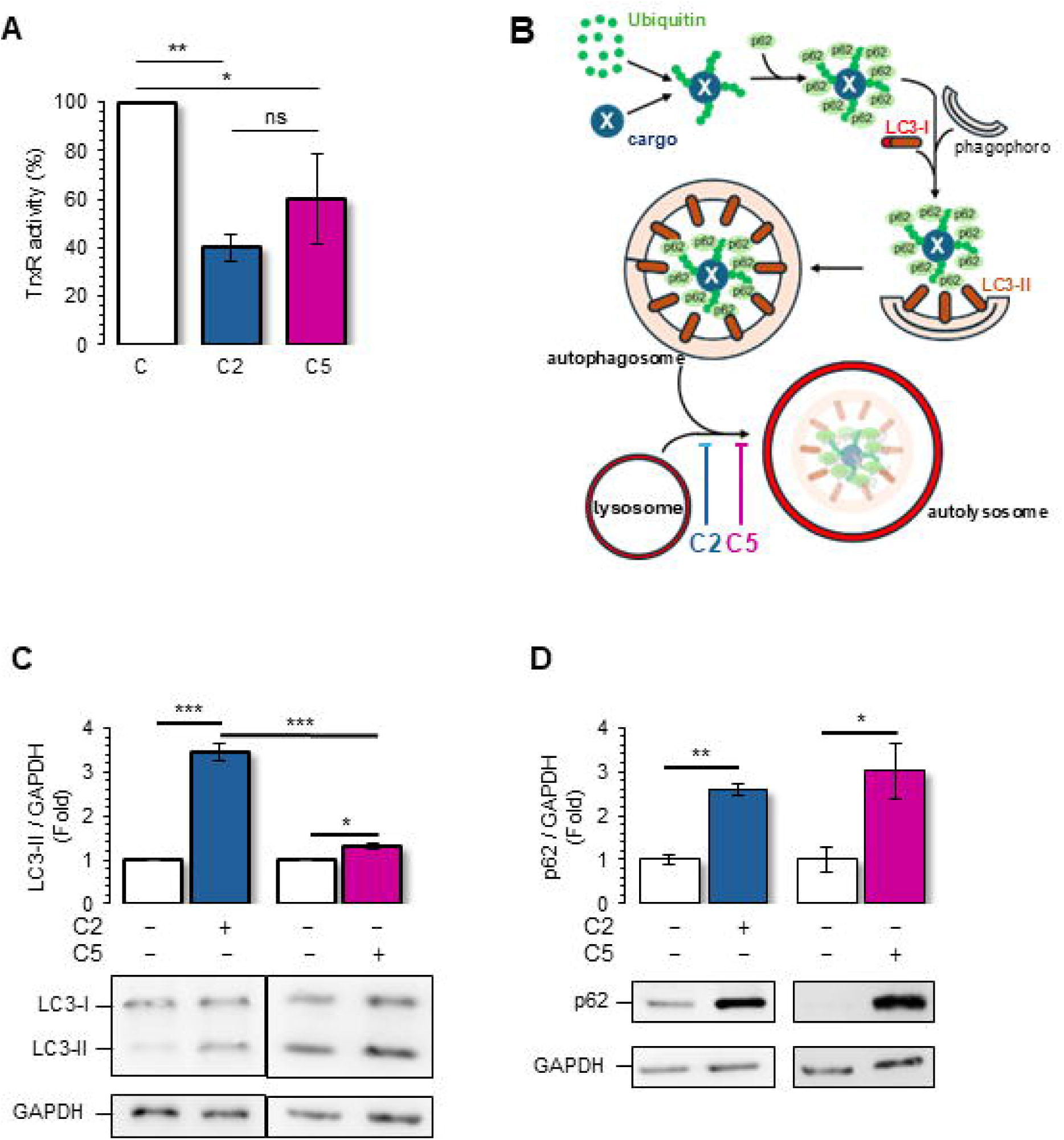
TrxR activity and autophagy pathway status. **A.** DMSO-treated cells (C) and cells treated with compounds **2** or **5** (C2 and C5) were collected 48 h after treatment, and TrxR activity was measured as described in the Materials and Methods section. **B.** Scheme of the autophagy pathway highlighting the roles of p62, LC3-I and -II and the proposed actions of compounds **2** and **5**. **C**. DMSO-treated cells (control) and cells treated with compounds **2** or **5** were collected 48 h after treatment, and the expression of LC3-II protein was evaluated by western blotting, as indicated in the Materials and Methods section. The LC3-II/GAPDH ratio is represented as a fold of difference from controls that were randomly assigned a value of 1. **D.** Same experiment as in C, but examining p62 levels relative to GAPDH. Quantification data are presented as mean ± SD of three independent experiments. Student’s t-test was used for statistical analysis. Statistical significance of data differences is indicated as follows: ns, no statistical significance; * p<0,05; ** p<0,01; *** p<0.001.

Finally, we analyzed the activation state of the autophagy pathway in cells in the presence and absence of these compounds. The two most used markers for autophagy are the phosphatidylethanolamine-conjugated forms of LC3-I (LC3-II) and SQSTM1/p62. When the autophagy pathway is activated in cells, LC3-II levels increase, whereas SQSTM1/p62 levels decrease (Figure 6B). The amount of LC3-II can be analyzed as the ratio of LC3-II to its precursor LC3-I, although some authors have reported that this method can yield inaccurate results because of the lack of sensitivity to LC3-I. Therefore, we chose to compare the ratio of LC3-II to GADPH, which generally presents more reliable and accurate results(43). The results showed that both compounds increased LC3-II levels (Figure 6C). This increase was larger for **C2** than for **C5** (Figure 6C), in agreement with the data on the inhibition of viral replication (Figures 2 and 4). Similar results were obtained when the LC3-II/LC3-I ratios were compared. The other autophagy marker used was SQSTM1/p62 (Figure 6B). When we compared the amount of this protein in the presence of **C2** and **C5**, we observed an increase in cells treated with these drugs compared with untreated cells (Figure 6D).

Therefore, we synthesized a series of compounds with improved biological properties, showing potent antiviral activity and promising selectivity indices against orthoflavivirus. Furthermore, we have been able to link the activity of these compounds to the modulation of cellular redox balance, suggesting interference with oxidative stress responses. These compounds may disrupt autophagy, thereby limiting viral propagation.

## DISCUSSION

The discovery and development of novel antiviral therapies remain challenging. In particular, orthoflaviviruses pose a serious threat to public health, as there are currently no available therapies or vaccines against them(19, 20). To address this gap, we systematically investigated a series of phosphaphenalene-gold(I) compounds for their effects on ZIKV, USUV, and BAGV replication. We verified their activity and specificity in terms of IC_50_ and SI, and their mechanism of action. Among these compounds, the most promising were **C2** and **C5,** which bear one and two gold-xanthate moieties, respectively (Table 1).

Comparison of infectious viral titers with intracellular viral RNA levels revealed a greater reduction in viral titers than in viral RNA copy numbers. This discrepancy may reflect methodological differences between the assays, as RT-qPCR detects viral RNA independently of infectivity, whereas infectious titration selectively measures replication-competent virus(44-46). Importantly, the more pronounced reduction observed in extracellular viral RNA suggests that the compounds could preferentially affect a late stage of the viral life cycle, whereas viral genome replication appears to be less strongly impacted. Although the precise mechanism remains unclear, alterations in the autophagy pathway may contribute to this effect. Autophagy is known to participate in intracellular membrane dynamics that are relevant for viral replication, assembly, and egress. The observed accumulation of LC3-II and p62 is consistent with a disruption of autophagic flux in cells treated with compounds **C2** and **C5**(47-50).

Phosphaphenalene gold(I) derivatives offer significant advantages in drug development because of their unique chemical properties and biological activities. One of their most notable features is the strong affinity of gold for thiol and selenol groups, allowing them to selectively interact with proteins such as TrxR(51). Their linear coordination geometry renders them synthetically accessible and tunable through ligands, such as those proposed in this study, which can improve stability, selectivity, and bioavailability in a simple manner(32, 33, 52).

A reduction in TrxR activity was observed in extracts from cells treated with these compounds, with **C2** showing a stronger effect than **C5** (Fig. 6A). This cellular enzyme is a target of other gold(I)-derived compounds(41). TrxR inhibition causes protein misfolding and activates clearance pathways, including autophagy, a key antiviral innate immune process(42, 53, 54). A basic analysis of autophagy involves the analysis of the levels of two proteins, LC3-II and SQSTM1/p62(43, 55). Generally, autophagy activation leads to the conversion of LC3-I to LC3-II with a consequent increase in the latter, while the p62 level decreases as it is degraded in the autolysosome (Figure 6B)(43). However, we observed an increase in both LC3-II and SQSTM1/p62 protein levels (Figures 6C and 6D). The simultaneous increase indicates that the cell is attempting to carry out autophagy (increased LC3-II), but there is an accumulation of material that is not being degraded properly (increased p62). This may be due to substrate overload, partial inhibition of autophagy, problems in the degradation of components within autophagosomes, or p62 transcriptional induction under redox stress(43, 55, 56). Similar to the gold(I) derivatives described in this study, bafilomycin A1, chloroquine and hydroxychloroquine induce autophagy with LC3-II accumulation while blocking autophagosome–lysosome fusion, leading to p62 accumulation(43, 57-60). The magnitude of TrxR inhibition (**C2**>**C5**) paralleled the antiviral potency and autophagy perturbation (Figures 2, 4, and 6), supporting a thioredoxin system-centered mechanism.

TrxR is a key enzyme in the thioredoxin system involved in cellular redox homeostasis and protection against oxidative stress(61). Viral infections can disrupt the balance between reactive oxygen species (ROS) and antioxidant defenses, leading to oxidative stress, as reported for SARS-CoV-2(62), HCV(63-65), HIV-1(65), influenza(66), chikungunya(67), and ZIKV(26, 27). Proteomic analysis of membranes from ZIKV-infected cells showed a significant amount of thioredoxin reductase 1 (TXNRD1)(68), the cytosolic form of TrxR. These authors proposed that TXNRD1 is recruited into membranes derived from the endoplasmic reticulum, where translation and replication of the viral genome occur, to ensure the correct folding of viral proteins containing disulfide bridges (e.g., E and NS1 proteins)(68). These data highlight the potential role played by TrxR. However, because DTNB-based measurements in lysates can be influenced by non-TrxR activities and residual compounds, our TrxR data should be interpreted as TrxR-associated rather than definitive.

It remains to be determined whether the inhibition of replication is due to the oxidation of viral proteins, host proteins necessary for replication, or both. Previous studies have indicated that viral proteins are directly affected by the oxidative state of the cell(23-25). Oxidation of viral proteins can compromise the ability of viruses to replicate. Enhancing the oxidation of viral proteins by various compounds (including Auranofin®) is a strategy that has also been effective in reducing the replication of chikungunya(67).

The potential of these compounds against other important RNA viruses, such as HIV, SARS CoV-2, and chikungunya, as well as other orthoflaviviruses, such as DENV and WNV, should be explored in the future. Furthermore, all antiviral and mechanistic data were generated in Vero cells, a robust flavivirus screening model. While these results provide in vitro proof-of-concept for the antiviral activity and mechanism of phosphaphenalene gold(I) complexes, validation in interferon-competent human models is required to confirm broader physiological relevance.

Overall, this study introduces a novel family of compounds, phosphaphenalene gold(I) complexes, with broad and remarkable therapeutic potential against ZIKV, USUV, and BAGV through the modulation of oxidative stress. Phosphaphenalenes stand out for their exceptional stability and unique chemical versatility, which allows precise fine-tuning of both biological and chemical properties. In summary, the phosphaphenalene gold(I) compounds exhibited low toxicity and high selectivity, highlighting their strong potential both as therapeutic candidates and as tools to advance the understanding of infection biology.

## Supporting information

Supplementary Informarion 1

## Acknowledgments

C.R.N. thanks the ERC for the Consolidator grant (ref: 101087685). Projects PID2021-125794OB-I00 and CNS2022-136028 funded by MICIU/AEI/10.13039/501100011033; PRTR-C17.I1 funded by MCIN with funding from the European Union NextGenerationEU and the JCCM; and SBPLY/21/180501/000185 and SBPLY/24/180225/000211 funded by JCCM and “ERDF A way to make Europe” are gratefully acknowledged. A.M. is supported by AEI funding with grant numbers PID2019-106068GB-I00 and PID2022-137974OB-I00 and JCCM funding with grant number SBPLY/21/180501/000076. We acknowledge the UCLM grant number 2022-GRIN-34150. We also acknowledge the technical assistance of Belén García Navarro.

## Declaration of generative AI and AI-assisted technologies in the manuscript preparation process

Pre-submission review was conducted using q.e.d. Science (https://www.qedscience.com). Paperpal was used to correct and refine the text.

