## Supplementary Informarion 1 for "Phosphaphenalene Gold(I) Complexes as Broad-Spectrum Antivirals Against Emerging Flaviruses"

[a] Department of Inorganic, Organic and Biochemistry, University of Castilla-La Mancha, Calle Almansa 14 - Edif. Bioincubadora, 02008, Albacete, Spain.
[b] Instituto Regional de Investigación Científica Aplicada (IRICA), University of Castilla-La Mancha, Avenida Camilo José Cela 1, 13071, Ciudad Real, Spain.
[c] Instituto de Investigación Biomédica, University of Castilla-La Mancha, Calle Almansa 14, 02008, Albacete, Spain.
[d] Unidad Asociada de Biomedicina UCLM-CSIC.
[e] Faculty of Pharmacy, University of Castilla-La Mancha, Avenida Dr. Jose Maria Sánchez Ibáñez, s/n 02008, Albacete, Spain.
[f] Faculty of Medicine, University of Castilla-La Mancha, Calle Almansa 14, 02008, Albacete, Spain.

† These authors contributed equally to the work.

1 Present address: mAbxience, Julia Morros s/n, 24009 León, Spain

### 1. General section

Reactions were carried out in dry glassware and under inert atmosphere of purified argon or nitrogen using Schlenk techniques. All solvents (THF, Toluene, DCM) were used directly from a solvent purification system MB SPS-800. Lithium diisopropylamine, P,P-diphenylchlorophosphine, trichlorosilane, chloro(dimethylsulfide)gold(I) and ehyl xhantogenate, were purchased from commercial suppliers and used as received.

**-NMR:** $^{1}H$, $^{13}C$, and $^{31}P$ NMR as well as COSY spectra were recorded on a 400 MHz Varian Inova NMR spectrometer. Chemical shifts are expressed as parts per million (ppm, δ) and referenced to external 85% $H_3PO_4$ ($^{31}P$), or solvent signals ($^{1}H$ / $^{13}C$): $CDCl_3$ (7.27 / 77.16 ppm) and $CD_2Cl_2$ (5.33 / 53.80 ppm) as internal standards. Signal descriptions include: s = singlet, d = doublet, t = triplet, q = quartet, m = multiplet and br = broad. All coupling constants are absolute values and J values are expressed in Hertz (Hz).

**-Mass spectrometry:** HR-ESI spectra were measured by the NUCLEUS analytical service of the University of Salamanca. GCMS was performed in a GC system 7250 GC/Q-TOF from Agilent Technologies.

### 2. Synthetic procedures

#### Compound 7

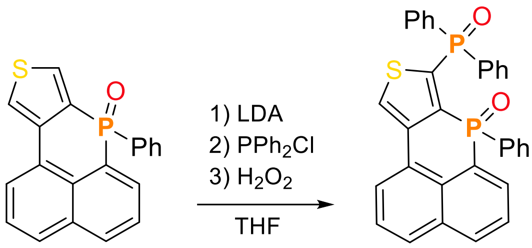


0.3 mL (1.1 eq, 0.132 mmol) of 1.0 M LDA was added to a solution of 7-phenyl-7H-benzo[4,5]phosphinolino[2,3-c]thiophene 7-oxide (1 eq, 0.120 mmol, 40 mg) dissolved in 2 ml of THF at -80°C. After the addition, the lithiated intermediate was allowed to react at room temperature. The mixture was stirred for 30 min at room temperature and 27 µL (1.2 eq, 0.144 mmol) of $PPh_2Cl$ at -80°C were added. The crude was stirred for 2 hours. After evaporation of the solvent under reduced pressure, the residue was redissolved in DCM. Subsequently, $H_2O$ and few drops of $H_2O_2$ were added. The mixture was stirred 15 min and the organic layer was separated. The latter was washed three times with water and dried over $Mg_2SO_4$. The product was purified by column chromatography using silica and AcOEt:EtOH 9.5:0.5 as eluent. Finally, 12 mg of brown color solid were obtained (19% yield).

**$^1$H NMR** (400 MHz, $CDCl_3$): δ ppm 8.44 (t, 1H), 8.25 (d, $J$ = 7.5 Hz, 1H), 8.14 (dd, $J$ = 14.4, 7.1 Hz, 1H), 8.06 – 7.98 (m, 3H), 7.89 (d, $J$ = 8.1 Hz, 1H), 7.66 – 7.51 (m, 5H), 7.43 (td, $J$ = 7.6, 3.1 Hz, 2H), 7.34 (t, $J$ = 7.3 Hz, 1H), 7.21 – 7.18 (m, 3H), 7.15 – 7.05 (m, 4H). **$^{13}$C{$^1$H} NMR** (101 MHz, $CDCl_3$): δ ppm 133.55, 133.47, 132.85, 132.74, 132.63, 132.40, 132.37, 132.34, 132.28, 132.21, 131.76, 131.11, 130.88, 130.72, 130.63, 129.86, 129.75, 129.64, 128.78, 128.66, 128.62, 128.47, 128.34, 128.30, 128.17, 128.07, 127.94, 126.65, 126.45, 126.32, 124.57. **$^{31}$P{$^1$H} NMR** (162 MHz, $CDCl_3$): δ ppm 23.0 (d, $J$ = 6.6 Hz), 6.5 (d, $J$ = 6.9 Hz). **HRMS** (ESI+) calcd. for $[M]^{\cdot+}$ $C_{32}H_{22}O_2P_2S^{\cdot+}$ 532.0816, found 532.0820.

#### Compound 8

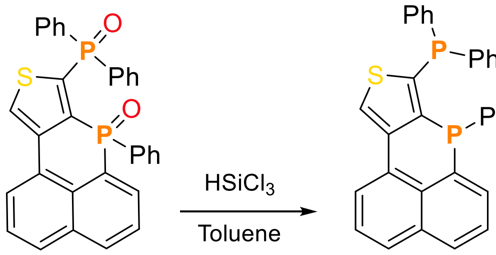


To a 15 ml Schlenk tube, dried and filled with nitrogen, compound **7** (1 eq, 0.056 mmol, 30 mg) was added and dissolved in 3 ml of dry toluene. Then, 0.114 mL of $HSiCl_3$ (20 eq, 1.12 mmol) were added dropwise at room temperature and the mixture was stirred at 100 ºC for two hours. The volatiles were removed under reduced pressure and the crude was filtered through aluminum oxide. 14.8 mg of a yellow solid were obtained (Yield: 52.8 %).

$^{1}$**H NMR** (400 MHz, $CDCl_3$): δ ppm 8.15 (s, 1H), 8.09 (d, *J* = 7.4, 1H), 7.86 – 7.81 (m, 3H), 7.54 (t, *J* = 7.6 Hz, 1H), 7.46 – 7.39 (m, 3H), 7.34 – 7.32 (m, 3H), 7.22 – 7.19 (m, 1H), 7.13 – 7.05 (m, 6H), 6.99 – 6.95 (m, 1H), 6.89 – 6.86 (m, 2H). $^{13}$**C{**$^{1}$**H} NMR** (101 MHz, $CDCl_3$): δ ppm 136.23, 136.13, 136.03, 135.53, 134.60, 134.43, 134.18, 133.48, 133.42, 133.29, 133.22, 132.93, 132.72, 132.05, 129.94, 129.72, 129.04, 128.90, 128.71, 128.58, 128.51, 128.36, 128.28, 128.19, 128.12, 127.93, 127.13, 126.21, 126.08, 125.94. $^{31}$**P{**$^{1}$**H} NMR** (162 MHz, $CDCl_3$): δ ppm -26.4 (d, *J* = 91.0 Hz), -46.2 (d, *J* = 91.2 Hz). **HRMS** (EI+) calcd. for $[M]^{\cdot+}$ $C_{32}H_{22}P_2S^{\cdot+}$ 500.0917, found 500.0924.

**Compound 9**

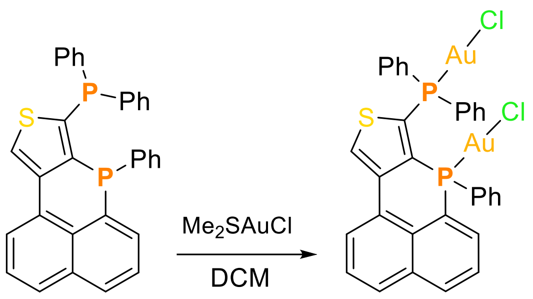


To a solution of compound **8** (1 eq, 0.03 mmol, 15 mg) in 2 ml of $CH_2Cl_2$, $Me_2SAuCl$ (2.05 eq, 0.06 mmol, 18.11 mg) was added at room temperature. The mixture was stirred 2 hours at room temperature and the volatiles were removed under vacuum. The crude was suspended in THF and precipitated used nPent. After washing with $Et_2O$, white solid was obtained (17.1 mg, yield: 60%).

$^{1}$**H NMR** (400 MHz, $CD_2Cl_2$): δ ppm 8.48 (s, 1H), 8.29 – 8.22 (m, 2H), 7.97 (dd, *J* = 22.0, 8.4 Hz, 2H), 7.70 – 7.57 (m, 5H), 7.55 – 7.46 (m, 5H), 7.43 – 7.29 (m, 5H), 7.19 (td, *J* = 7.5, 2.6 Hz, 2H). $^{13}$**C {**$^{1}$**H} NMR** (101 MHz, $CD_2Cl_2$): δ ppm 137.96, 137.73, 137.45, 137.21, 135.05, 134.90, 134.88, 134.73, 134.52, 133.98, 133.95, 133.34, 133.29, 133.26, 133.16, 133.01, 132.32, 132.30, 131.08, 130.14, 130.12, 130.01, 129.82, 129.74, 129.69, 128.79, 128.47, 127.82, 127.49, 126.83, 126.66.$^{31}$**P{**$^{1}$**H} NMR** (162 MHz, $CD_2Cl_2$): δ ppm 15.2 (d, *J* = 40.8 Hz), -0.8 (d, *J* = 40.8 Hz). **HRMS** (ESI+) calcd. for ($[M – Cl]^{\cdot+}$) $C_{32}H_{22}Au_2ClP_2S$ 928.9937, found 928.9916.

**Compound 5**

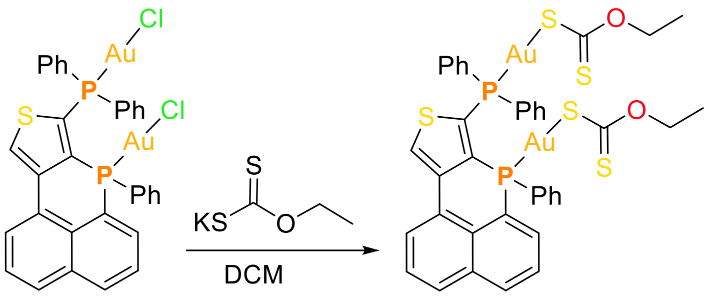


In the darkness, compound **9** (1 eq, 0.016 mmol, 16 mg) was dissolved in 1 ml of DCM and potassium ethyl xanthogenate (1 eq, 0.033 mmol, 5.56 mg) was added at room temperature. Then, the mixture was stirred for 3 hours at room temperature and the volatiles were removed under vacuum. The crude was redissolved in DCM and $H_2O$ was added. The organic layer was separated and dried over $Mg_2SO_4$.The product was washed with n-pentane. Green-Yellow solid was obtained (17.1 mg, yield: 40%).

**$^1$H NMR** (400 MHz, $CD_2Cl_2$): δ ppm 8.51 (s, 1H), 8.40 – 8.27 (m, 2H), 7.94 (dd, $J$ = 19.5, 8.2 Hz, 2H), 8.74 –8.60 (m, 3H), 7.60 – 7.44 (m, 7H), 7.38 (dd, $J$ = 13.7, 7.1 Hz, 2H), 7.30 – 7.22 (m, 3H), 7.11 (td, $J$ = 7.7, 2.8 Hz, 2H), 4.45 (q, $J$ = 7.1 Hz, 4H), 1.33 (t, $J$ = 7.1 Hz, 6H). **$^{13}$C {$^1$H} NMR** (101 MHz, $CD_2Cl_2$): δ ppm 225.10, 137.17, 136.93, 135.08, 134.93, 134.82, 134.67, 134.49, 134.43, 134.10, 133.60, 133.57, 133.53, 133.04, 133.02, 132.95, 132.80, 132.25, 131.92, 131.89, 130.90, 129.99, 129.86, 129.71, 129.69, 129.59, 129.57, 127.50, 127.44, 126.84, 126.67, 125.29, 70.64, 14.53.**$^{31}$P{$^1$H} NMR** (162 MHz, $CD_2Cl_2$): δ ppm 19.8 (d, $J$ = 40.8 Hz), 1.5 (d, $J$ = 40.6 Hz). **HRMS** (ESI+) calcd. for ($[M – C_3H_5OS_2]^{\cdot+}$) $C_{35}H_{27}Au_2OP_2S_3$ 1015.0030, found 1015.0004.

### 3. NMR data

#### [1]H NMR (400 MHz, CDCl$_3$) of Compound 7

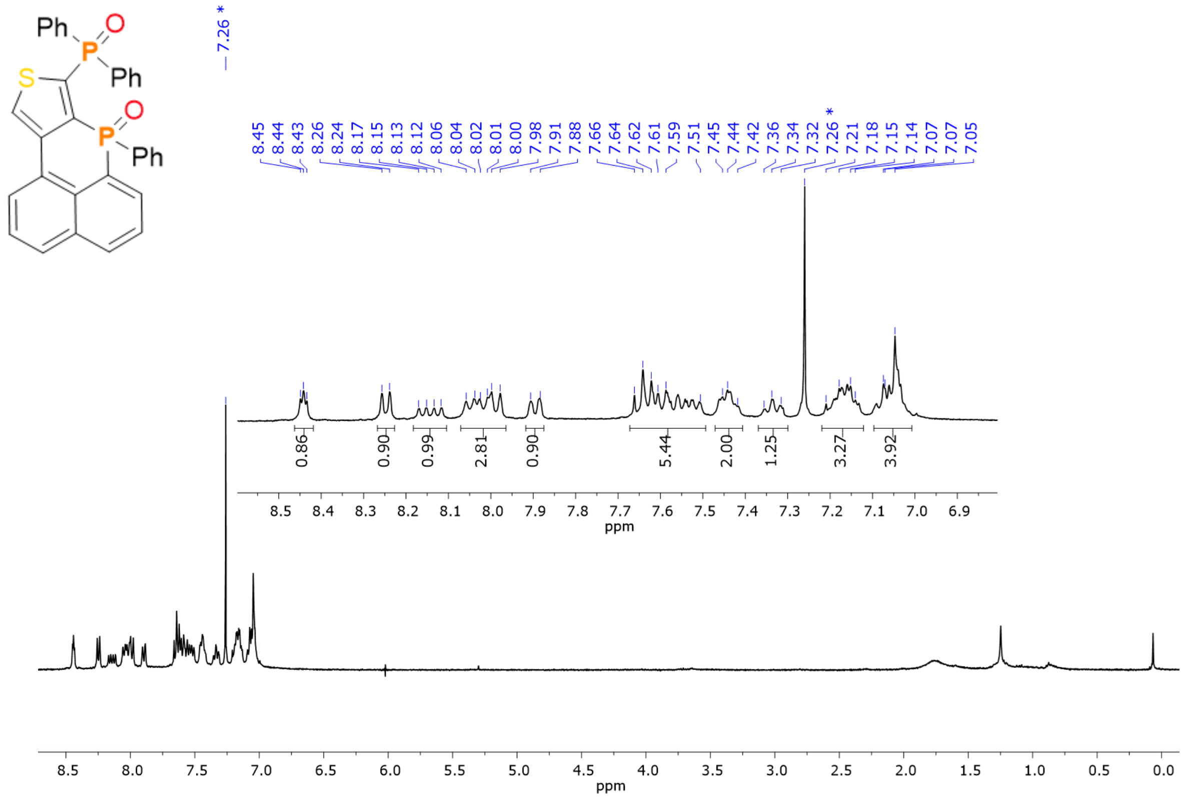

#### [13]C{[1]H} NMR (400 MHz, CDCl$_3$) of Compound 7

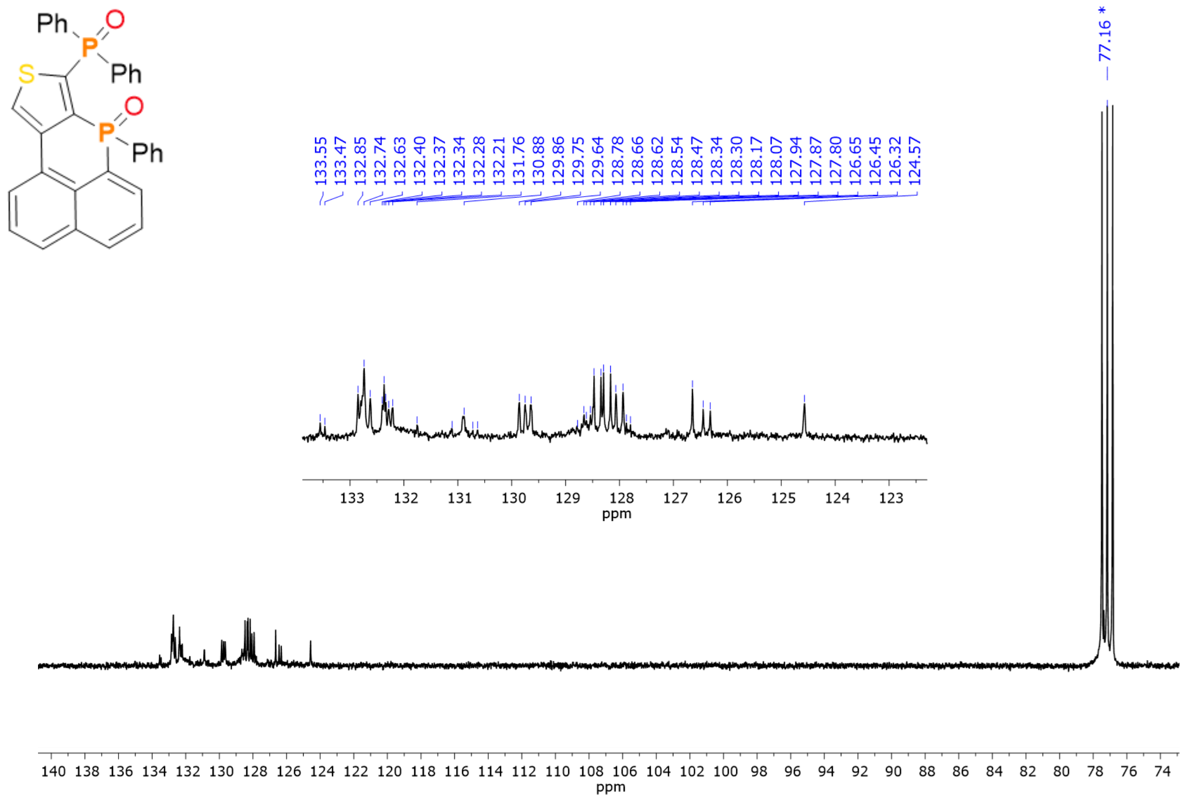

**$^1$H-$^1$H COSY NMR (400 MHz, $CDCl_3$) of Compound 7**

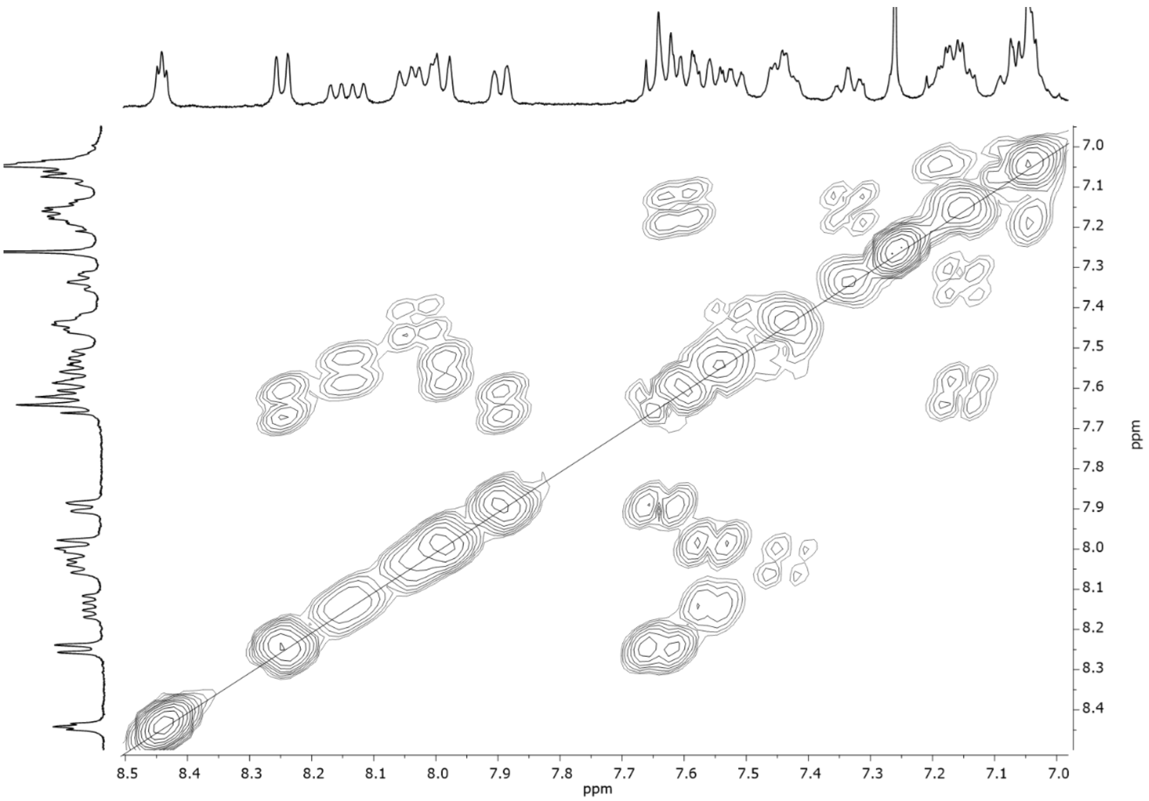


**$^{31}$P{$^1$H} NMR (400 MHz, $CDCl_3$) of Compound 7**

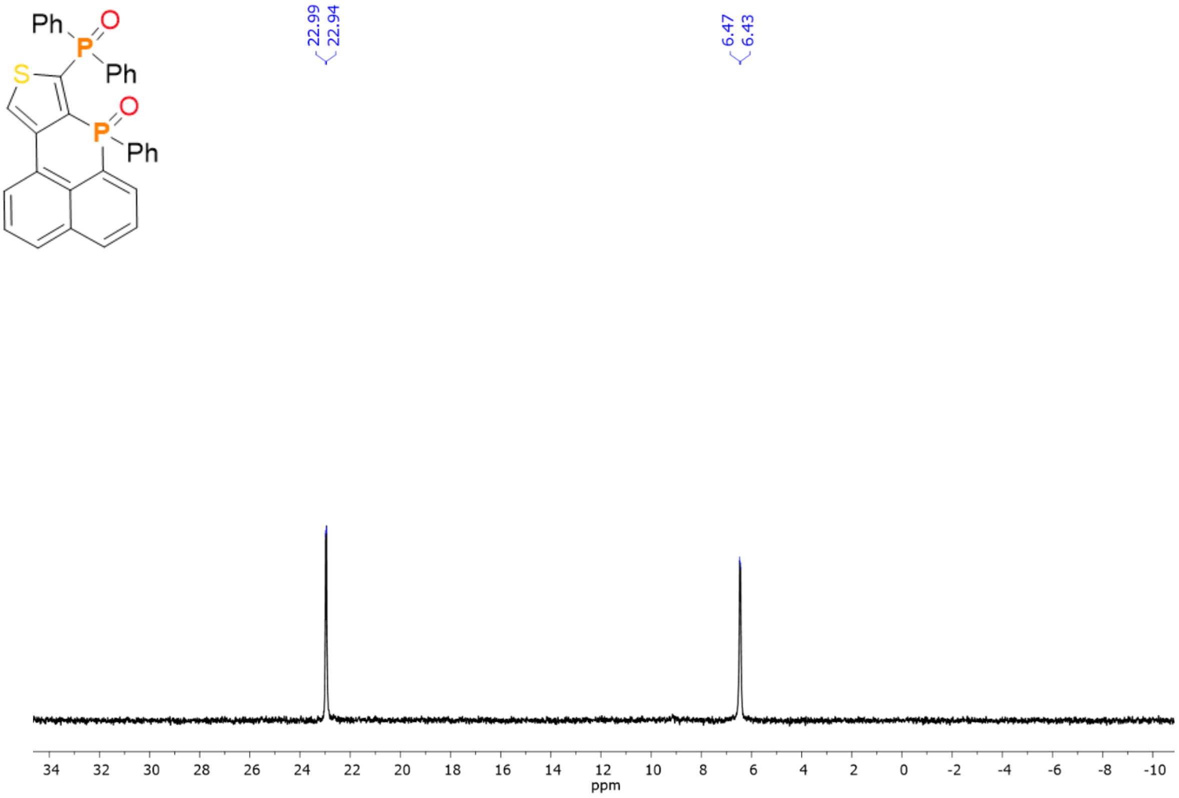

[1]H NMR (400 MHz, $CDCl_3$) of Compound 8

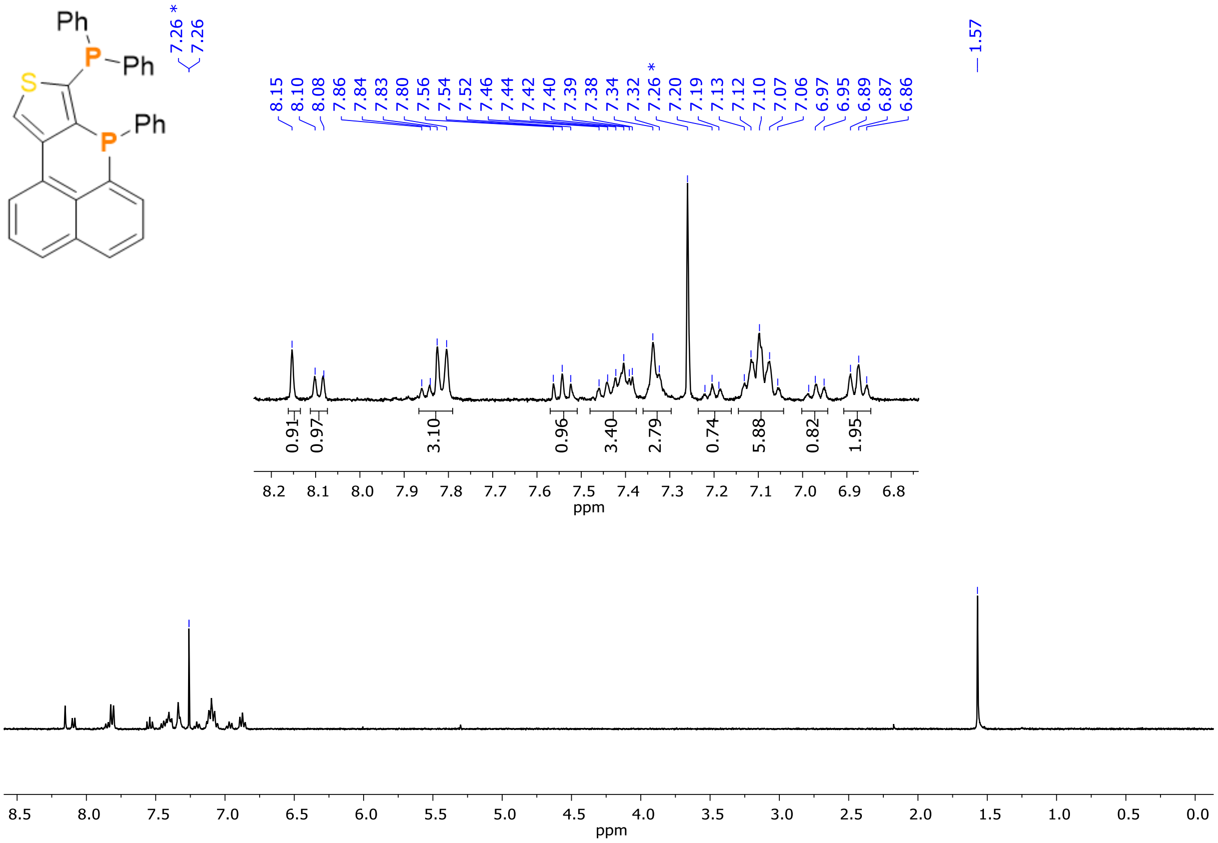


[13]C{[1]H} NMR (400 MHz, $CDCl_3$) of Compound 8

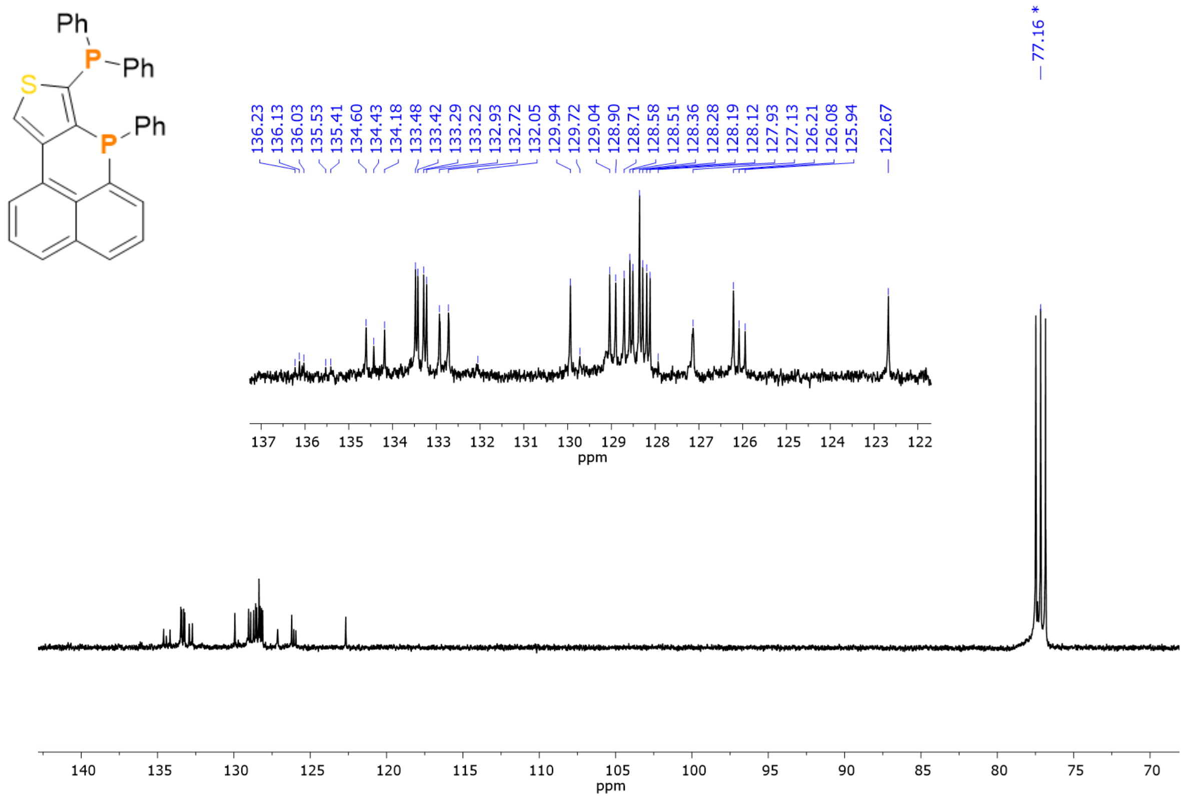

**$^{1}H$-$^{1}H$ COSY NMR (400 MHz, $CDCl_3$) of Compound 8**

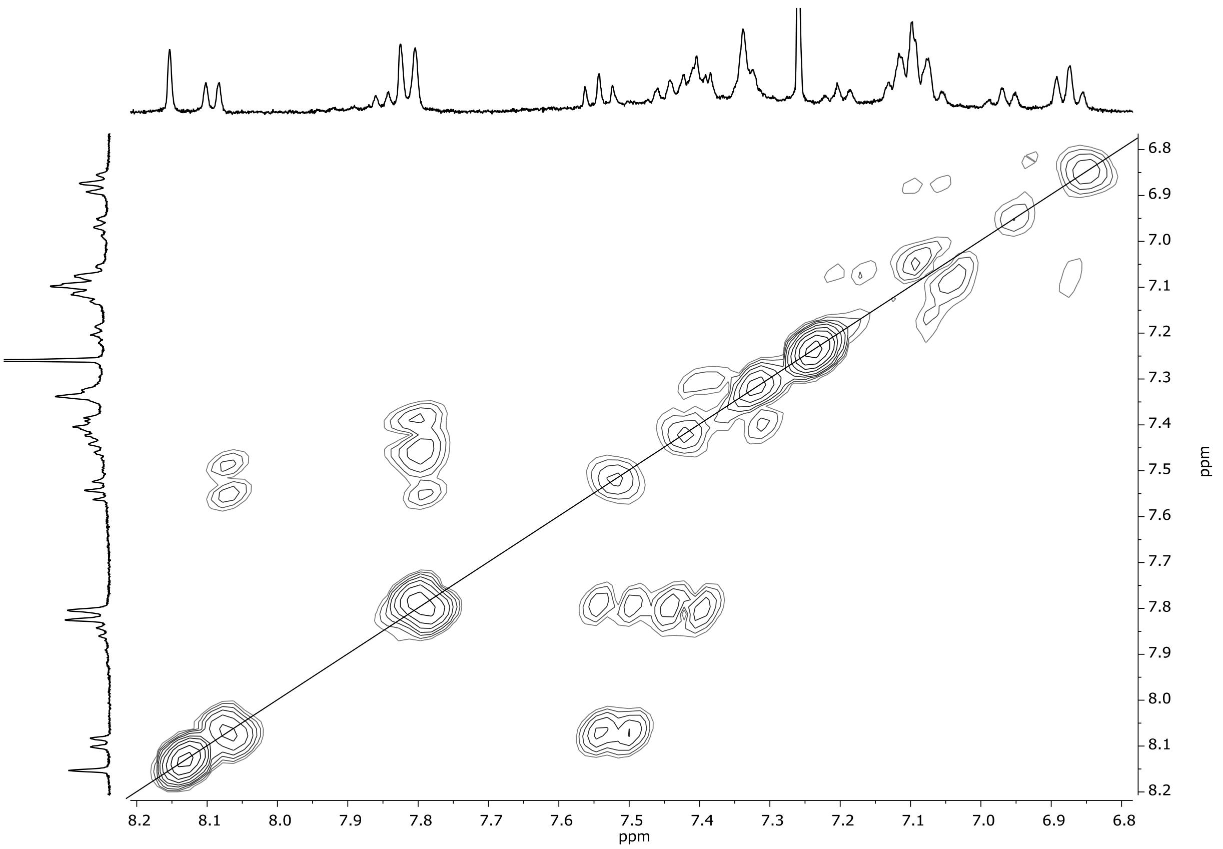


**$^{31}P\{^{1}H\}$ NMR (400 MHz, $CDCl_3$) of Compound 8**

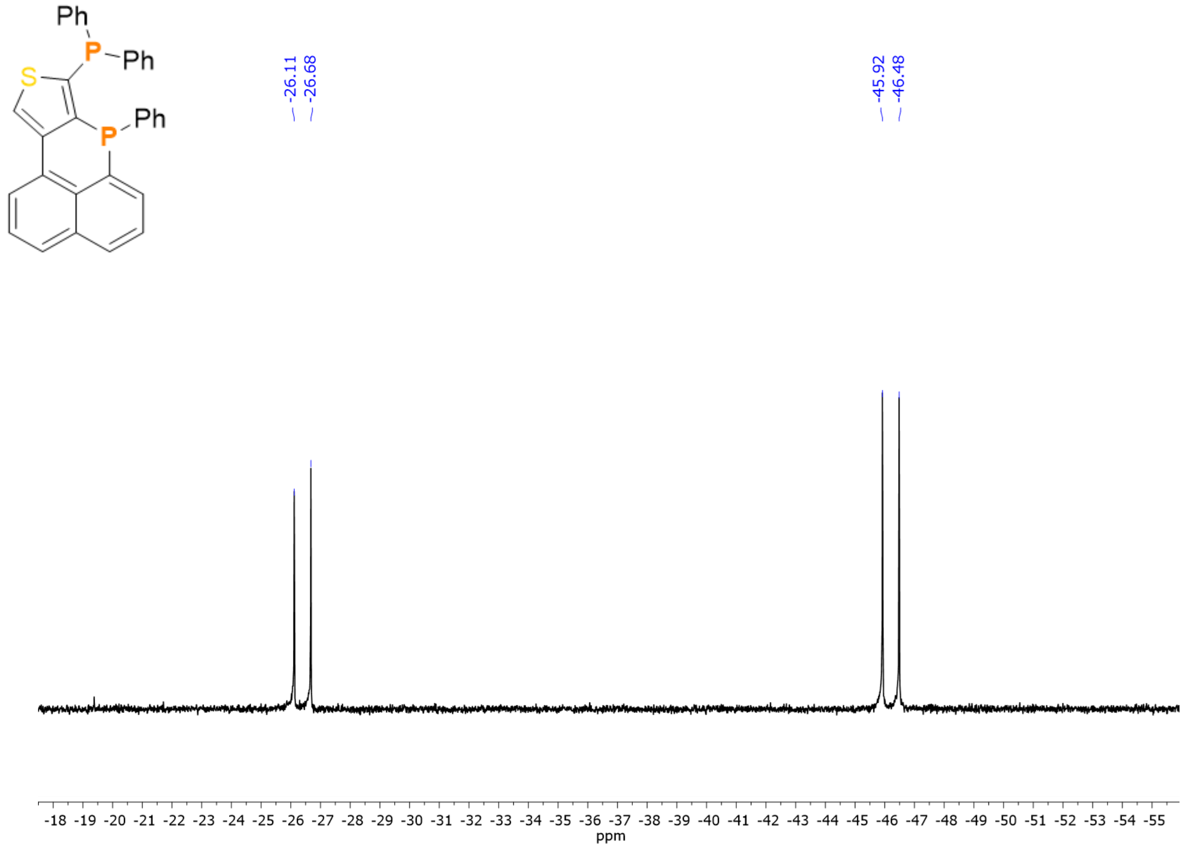

$^{1}H$ NMR (400 MHz, $CD_2Cl_2$) of Compound 9

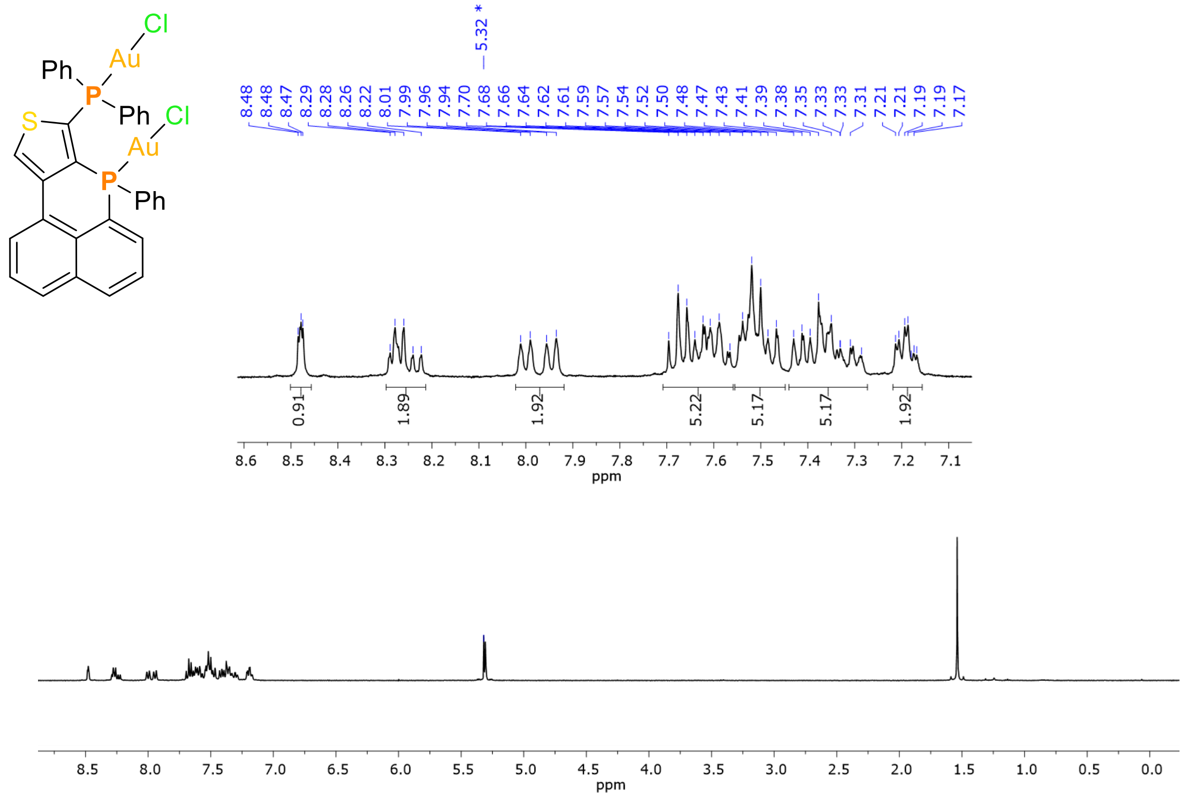


$^{13}C\{^{1}H\}$ NMR (400 MHz, $CD_2Cl_2$) of Compound 9

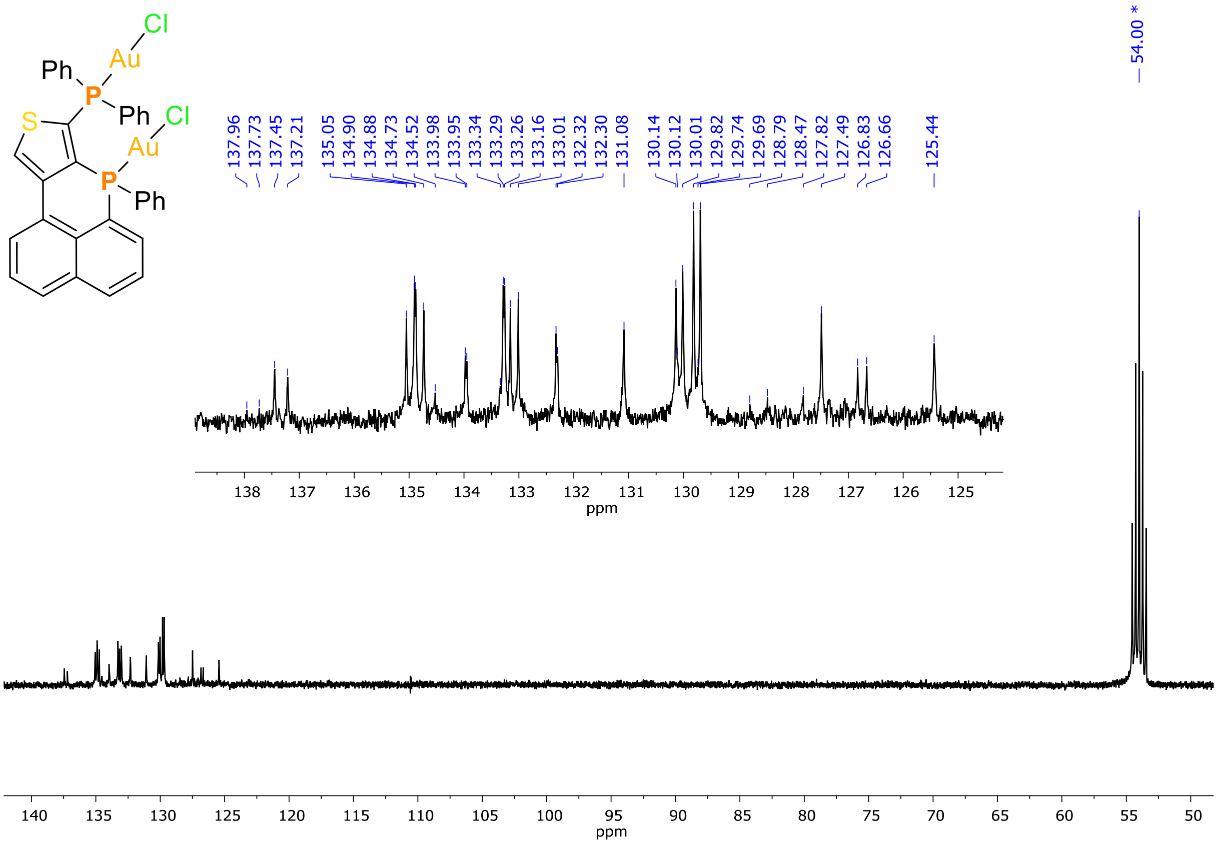

$^{1}H$-$^{1}H$ COSY NMR (400 MHz, $CD_2Cl_2$) of Compound 9

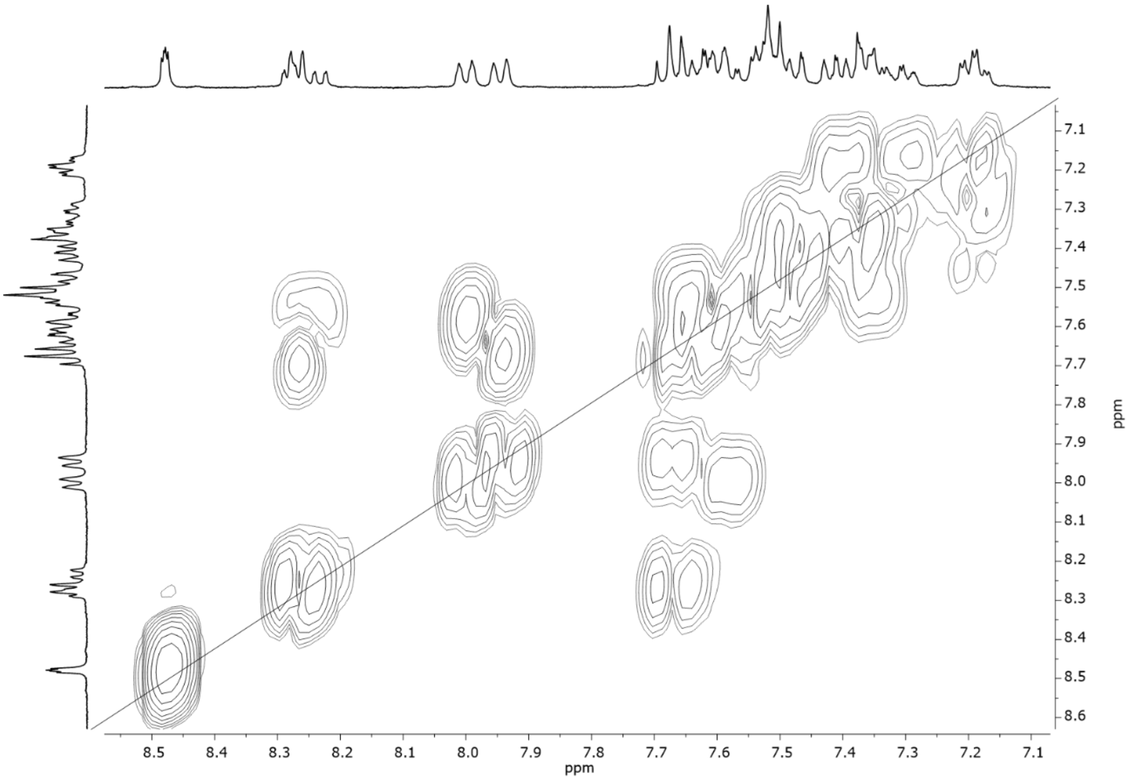


$^{31}P\{^{1}H\}$ NMR (400 MHz, $CD_2Cl_2$) of Compound 9

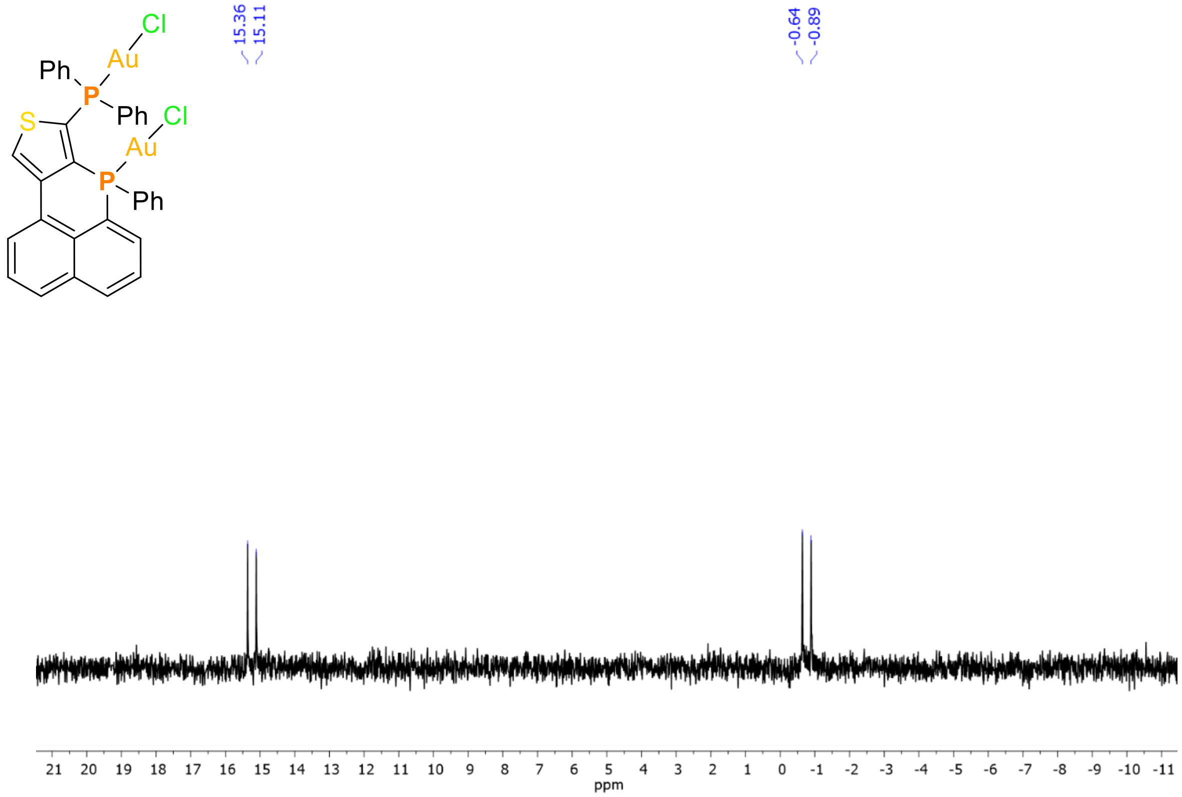

**$^{1}$H NMR (400 MHz, $CD_2Cl_2$) of Compound 5**

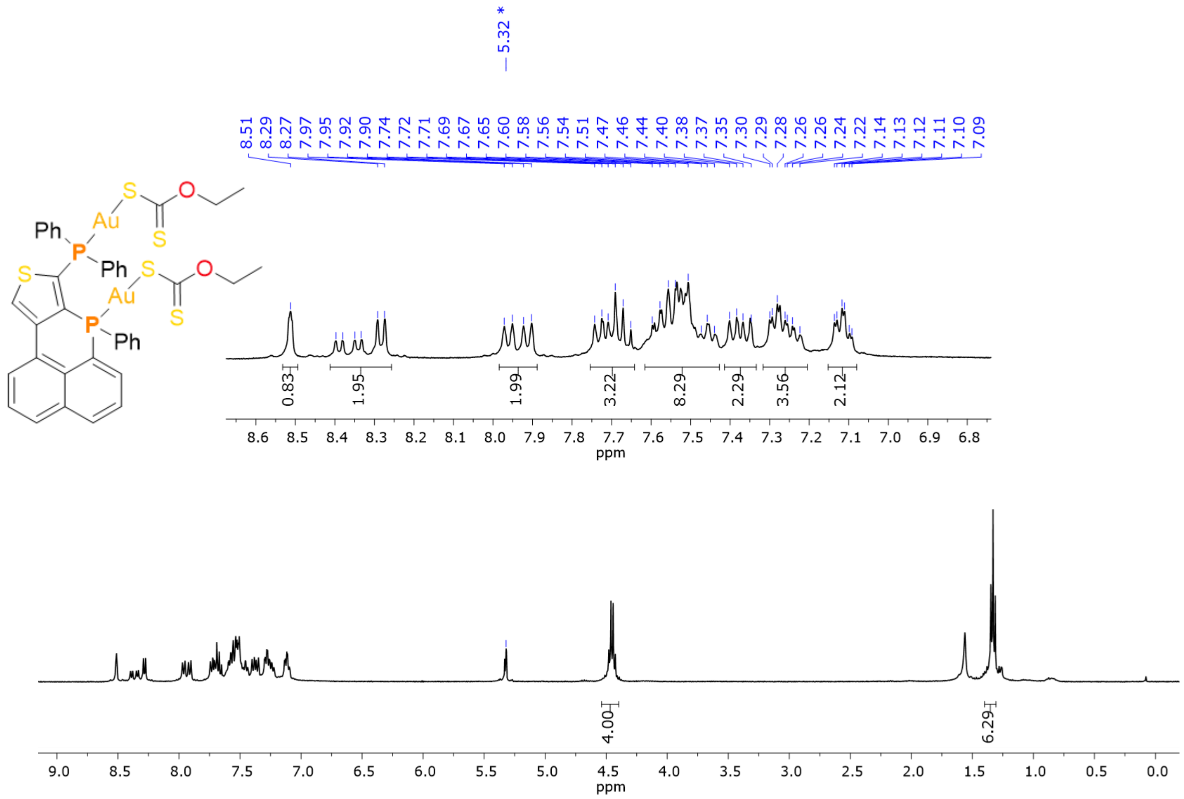


**$^{13}$C{$^{1}$H} NMR (400 MHz, $CD_2Cl_2$) of Compound 5**

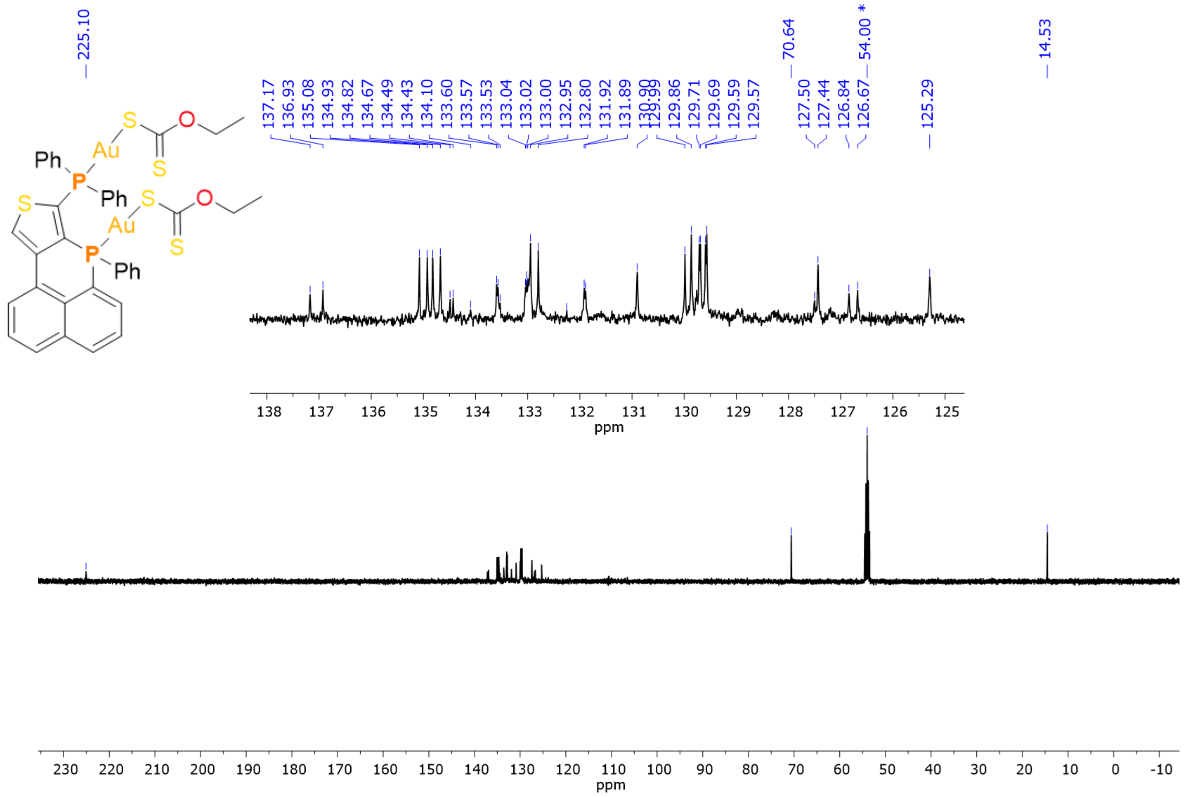

$^{1}H$-$^{1}H$ COSY NMR (400 MHz, $CD_2Cl_2$) of Compound 5

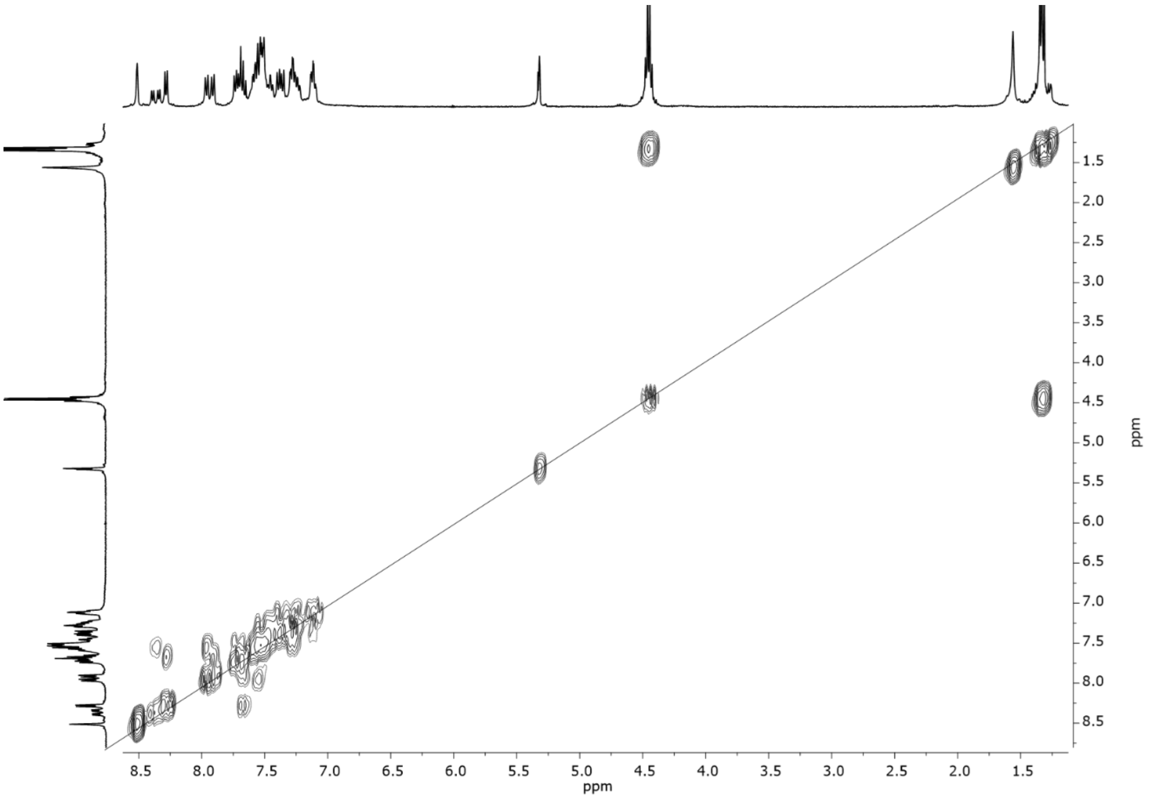

$^{31}P\{^{1}H\}$ NMR (400 MHz, $CD_2Cl_2$) of Compound 5

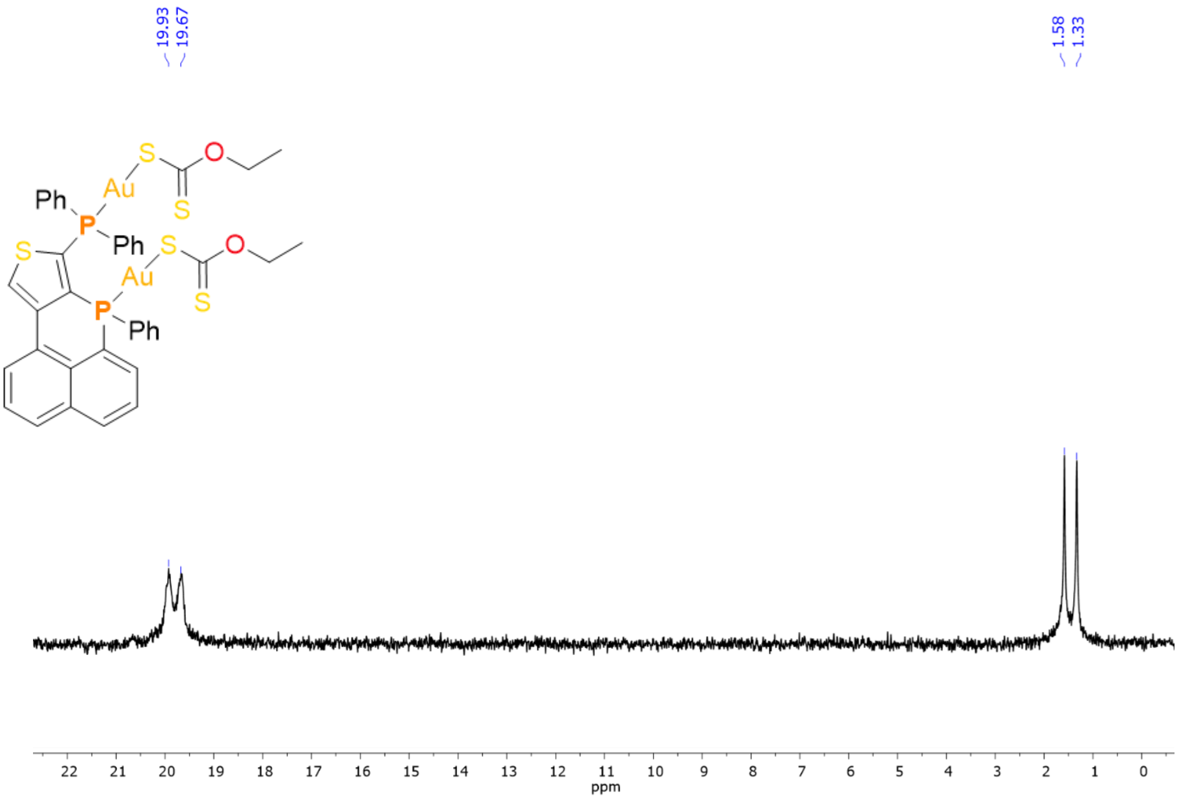
